# Neural dynamics in the limbic system during male social behaviors

**DOI:** 10.1101/2023.03.12.532199

**Authors:** Zhichao Guo, Luping Yin, Takuya Osakada, Julieta Lischinsky, Jonathan Chien, Bing Dai, Ashley Urtecho, Xiaoyu Tong, Zhe S. Chen, Dayu Lin

## Abstract

Sexual and aggressive behaviors are two evolutionarily conserved social behaviors vital for an animal’s survival and reproductive success. While an increasing number of brain regions in the limbic system have been identified as functionally relevant for these two types of behaviors, an understanding of how social cues are represented across brain regions and how social behaviors are generated via this network activity remains elusive. To gain a holistic view of the neural responses during social behaviors, we utilized multi-fiber photometry to simultaneously record Ca^2+^ signals of estrogen receptor alpha *(Esr1)*-expressing cells from 13 limbic brain regions in male mice during sexual and aggressive behaviors and compare the response magnitude and temporal patterns across regions. We find that conspecific sensory information, as well as social action initiation signals, are widely distributed in the limbic system and can be decoded from the network activity. Cross-region correlation analysis reveals striking increases in functional connectivity in the network during the action initiation phase of social behaviors whereas advanced copulation is accompanied by a “dissociated” network state. Based on the response patterns, we propose a mating-biased network (MBN) and an aggression-biased network (ABN) for mediating male sexual and aggressive behaviors, respectively.

## Introduction

Sexual and aggressive behaviors are two fundamental social behaviors. For males, sexual reproduction entails properly displaying copulative behaviors toward conspecific females, while the ability to deploy aggression to fend off conspecific male intruders is key to securing resources for mating success. These behaviors are innate, i.e., inborn, and thus, their generation should be supported by developmentally wired circuits. In 1999, Sarah Newman proposed the existence of a social behavior network (SBN) that mediates innate social behaviors in mammals based on decades of lesion and immediate early gene mapping studies (Newman, 1999). The SBN includes seven interconnected subcortical areas: medial amygdala (MeA), bed nucleus of stria terminalis (BNST), medial preoptic area (MPOA), anterior hypothalamus (AHN), lateral septum (LS), ventromedial hypothalamus (VMH), and midbrain (including periaqueductal gray (PAG) and tegmentum) (Newman, 1999). MeA and BNST are collectively called the extended medial amygdala. In 2005, James Goodson extended this network to non-mammalian vertebrate species based on studies in birds and teleost (bony) fish (Goodson, 2005). In recent years, the importance of SBN in social behaviors has been continuously validated and elaborated. For instance, gain- and loss-of-function studies demonstrated an indispensable role of the ventrolateral part of the ventromedial hypothalamic nucleus (VMHvl) in aggressive behaviors in mice (Falkner et al., 2016; Hashikawa et al., 2017; Lee et al., 2014; Lin et al., 2011; Yang et al., 2013; Yang et al., 2017), while the molecular identities of cells in the medial preoptic nucleus (MPN) relevant for sexual behaviors have been increasingly refined (Gao et al., 2019; Karigo et al., 2020; Michael et al., 2020; Wei et al., 2018).

A basic feature of the SBN is its enrichment of gonadal hormone receptors, which allow the cells to be modulated by gonadal steroid hormones, including androgens (testosterone), estrogens, and progesterone (Newman, 1999). Indeed, gonadal hormones are crucial for the emergence of social behaviors during development and their maintenance during adulthood in both males and females (Jennings and de Lecea, 2020; Wu and Shah, 2011). For example, female mice exposed to prenatal testosterone show male-like sexual and aggressive behaviors during adulthood (Edwards and Burge, 1971). Adult castration abolishes masculine behaviors, which can be restored by testosterone supplements (McCarthy, 2008). In males, testosterone mainly acts through estrogen receptors after being converted into estrogen via the enzyme aromatase (Wu and Shah, 2011). Knocking out estrogen receptor alpha (Esr1) disrupts male sexual and aggressive behaviors severely (Ogawa et al., 2000; Ogawa et al., 1997; Wersinger et al., 1997). Thus, it is perhaps not a coincidence that Esr1-expressing cells in the MPN and VMHvl are found to be the relevant populations for sexual and aggressive behaviors (Hashikawa et al., 2017; Karigo et al., 2020; Lee et al., 2014; Wei et al., 2018).

The SBN does not cover all regions essential for social behaviors and requires an expansion. Several regions outside the SBN have recently been identified as necessary for male sexual or aggressive behaviors, or both. Posterior amygdala (PA) cells promote aggression and sexual behaviors through projections to VMHvl and MPN, respectively (Stagkourakis et al., 2020; Yamaguchi et al., 2020; Zha et al., 2020). The ventral part of the premammillary nucleus (PMv), a hypothalamic nucleus posterior to the VMHvl, projects heavily to both MPN and VMHvl and is important for male aggression (Chen et al., 2020; Soden et al., 2016; Stagkourakis et al., 2018). The ventral subiculum has also been found to bi-directionally modulate aggression at least partly through its projection to VMHvl (Chang and Gean, 2019). Interestingly, these newly identified social behavior-relevant regions share the same features of the original SBN: high levels of sex hormone receptor expression and extensive connections with regions in the SBN (Canteras et al., 1992a, b; Mitra et al., 2003; Yamaguchi et al., 2020).

Every behavior is a phenotypical manifestation of some well-orchestrated network activity. As our knowledge of the functions of individual brain regions and connections in social behaviors accumulates, an important next step is to holistically understand the behavior-relevant neural activity in a large network of interacting regions. Indeed, several recent studies have attempted to achieve this goal in other behavioral contexts using large-scale single-unit recording with multi-site silicon probes, e.g., Neuropixels (Allen et al., 2019; Juavinett et al., 2019; Jun et al., 2017; Siegel et al., 2015; Steinmetz et al., 2019). However, recording sites of silicon probes are generally distributed along the vertical shafts, making them unsuitable for simultaneous recording from multiple deep subcortical regions. As an alternative approach, fiber photometry, a method first developed to record bulk fluorescence signals from subcortical regions (Cui et al., 2013; Gunaydin et al., 2014), has also been scaled up to enable recording from multiple regions (Guo et al., 2015; Kim et al., 2016). Here, the lack of cellular resolution is offset by the ability to record from molecularly-defined subpopulations, or in other words, cells with potentially similar functions and relatively homogeneous responses (Lin and Schnitzer, 2016). The recent incorporation of high-density customizable multi-fiber arrays dramatically reduces the total weight of the implant and associated brain damage, making it well-suited for recording multiple sites in the SBN and beyond in freely moving animals (Sych et al., 2019).

Leveraging this multi-fiber photometry (MFP) technique, we simultaneously recorded the Ca^2+^ activities of *Esr1*^+^ populations from 13 regions in the limbic system, referred to here as the expanded SBN, during sexual and aggressive behaviors in freely-moving male mice. Our results reveal dynamic neural representations of these two types of behaviors at the network level.

## Results

### Simultaneous recording from 13 brain regions in the extended social behavior network

We selected 13 brain regions that have either been implicated in aggressive and/or sexual behaviors or are strongly connected with those regions for Ca^2+^ recording during social behaviors. The list includes five hypothalamic regions – medial preoptic nucleus (MPN), anterior hypothalamic nucleus (AHN), ventrolateral part of the ventromedial hypothalamus (VMHvl), dorsomedial hypothalamus (DMH) and ventral premammillary nucleus (PMv), five amygdala regions –anterior medial amygdala (MeAa), posterodorsal medial amygdala (MeApd), posterior amygdala (PA), posteromedial cortical amygdala (CoApm) and posteromedial bed nucleus of stria terminalis (BNSTpm), and three regions outside of amygdala and hypothalamus – ventral part of lateral septum (LSv), ventral subiculum (SUBv) and lateral periaqueductal gray (lPAG) (**Figure 1A**) (Bayless et al., 2019; Chang and Gean, 2019; Chen et al., 2020; Falkner et al., 2020; Hong et al., 2014; Karigo et al., 2020; Lee et al., 2014; Lenschow and Lima, 2020; Leroy et al., 2018; Lin et al., 2011; Lischinsky and Lin, 2020; Newman, 1999; Stagkourakis et al., 2020; Stagkourakis et al., 2018; Unger et al., 2015; Wei et al., 2018; Wong et al., 2016; Xie et al., 2020; Yamaguchi et al., 2020; Yang et al., 2013; Yang et al., 2017; Zelikowsky et al., 2018; Zha et al., 2020; Zhu et al., 2020). Each of these 13 regions is densely connected with multiple other recorded regions and expresses abundant *Esr1,* with the exceptions of AHN, LSv, and PAG, where *Esr1* expression is modest. To target *Esr1*-expressing cells in these regions, we injected Cre-dependent GCaMP6f viruses into each candidate region in Esr1-2A-Cre male mice (**Figures S1 and S2**). During the same surgery, two custom arrays, each composed of multiple 100-µm optic fibers, with one targeting seven medially located regions and the other targeting five laterally positioned regions, and a single fiber targeting lPAG were implanted (**Figures 1B**). The recording setup is a modified version of previously reported setups (Kim et al., 2016; Sych et al., 2019). It uses a low-cost CCD camera to capture images from the end of a 19-channel fiber bundle (only 13 channels were used here) that delivers and collects, respectively, the excitation and emission light from each recording site (**Figures 1C, D, E**).

**Figure 1:**
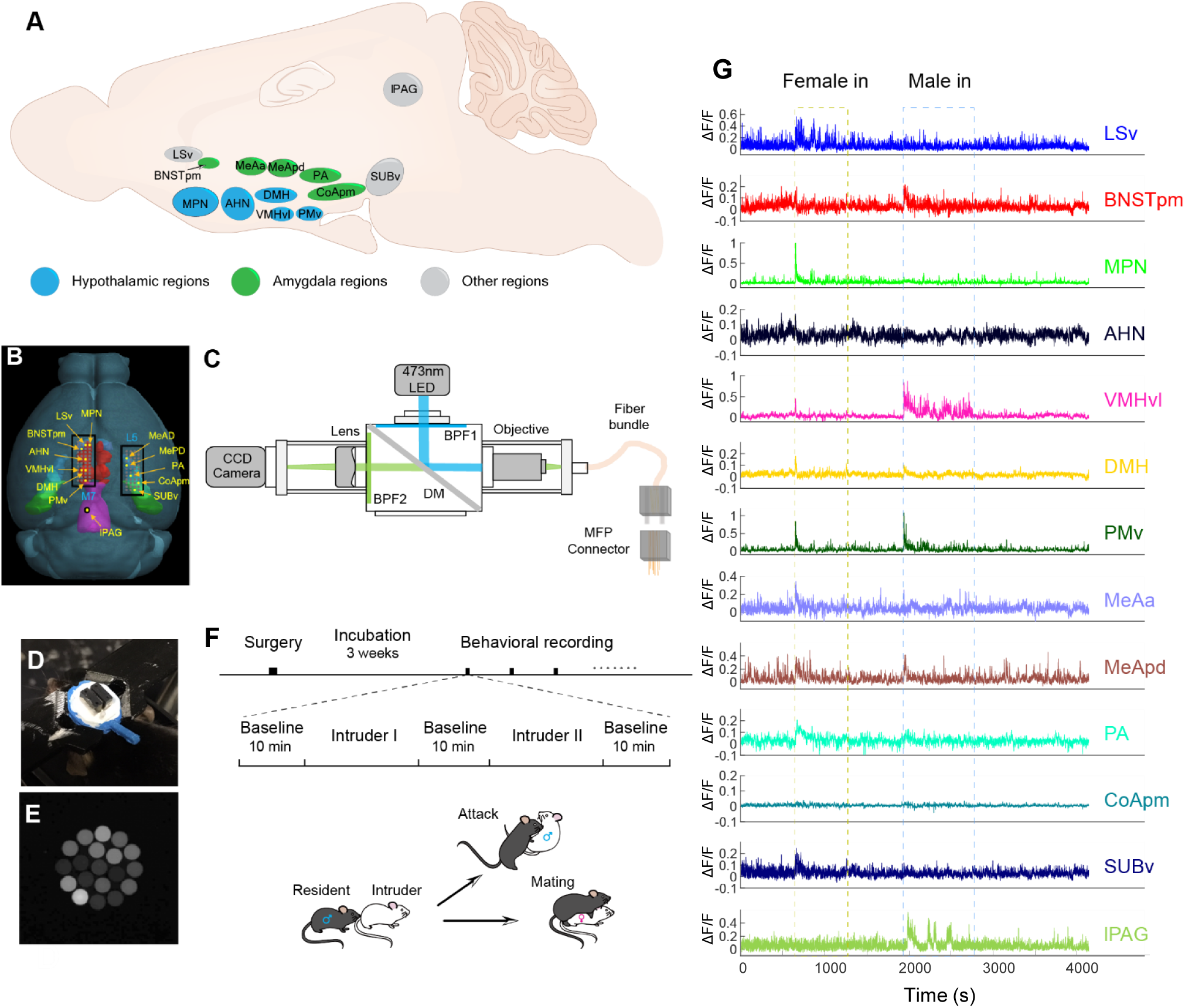
Multi-fiber photometry (MFP) recording of 13 regions in the limbic system. **A.** Illustration showing the recorded regions in mouse hypothalamus (blue), amygdala (green), and other brain areas (gray). **B.** The optic fiber arrays overlaid on a 3D mouse brain model showing various targeted structures. The model is from https://connectivity.brain-map.org/. **C.** Diagram of the MFP recording system. **D.** An animal with the implanted fiber arrays and a head-fixation ring. **E.** An image showing the end of the optic fiber bundle. **F.** Experimental and each recording session timeline. **G.** Simultaneously recorded GCaMP6f traces (ΔF/F) from a representative recording session.

Recordings started three weeks after the surgery, each comprised of a male, a female, and sometimes an object session (**Figure 1F**). During the male session, a group-housed non-aggressive Balb/C male was introduced into the home cage of the recording male for approximately 10 minutes, while a receptive female was introduced during the female session. We observed peak fluorescence changes (ΔF/F) over 100%, comparable to the Esr1 cell responses recorded using a conventional single-fiber photometric recording setup (Bayless et al., 2019; Chen et al., 2020; Falkner et al., 2016; Falkner et al., 2020; Wang et al., 2019; Wei et al., 2018; Yamaguchi et al., 2020) (**Figure 1G**). We performed the recording two to four times for each animal, with 3-7 days between recording sessions. Histological analysis was performed for all animals, and only correctly targeted brain regions were used for a given animal. The final dataset contains 64 recording sessions from 25 animals (mean ± STD = 2.6 ± 1.0 sessions/animal), with 10.6 ± 2.2 (mean ± STD) recording sites/animal.

### Large activity increase across the expanded SBN during the initial encounter with a conspecific

Upon entry of an intruder, regardless of sex, many brain regions in the male mice showed remarkable increases in Ca^2+^ activity (**Figures 2A-F**). During the male introduction, VMHvl showed the largest activity increase among all recorded regions, followed by DMH and PMv (**Figure 2A, E, G**, **and** H). During the female introduction, MPN showed the largest increase, with VMHvl, DMH, PMv, MeAa, BNSTp, MeAp, and PA all showing a similarly large increase (**Figure 2I, J**). Outside the hypothalamus and amygdala, SUBv and LSv also increased activity moderately, whereas the activity increase in the lPAG was minor and only significant during the male introduction (**Figure 2G-J**). When comparing the entry response towards males and females, VMHvl, DMH, and PMv showed a preferential response towards males over females, whereas MPN, BNSTp, MeAa, and PA showed the opposite response bias, suggesting their potentially preferential involvement in male- or female-directed social behaviors (**Figure 2S**).

**Figure 2:**
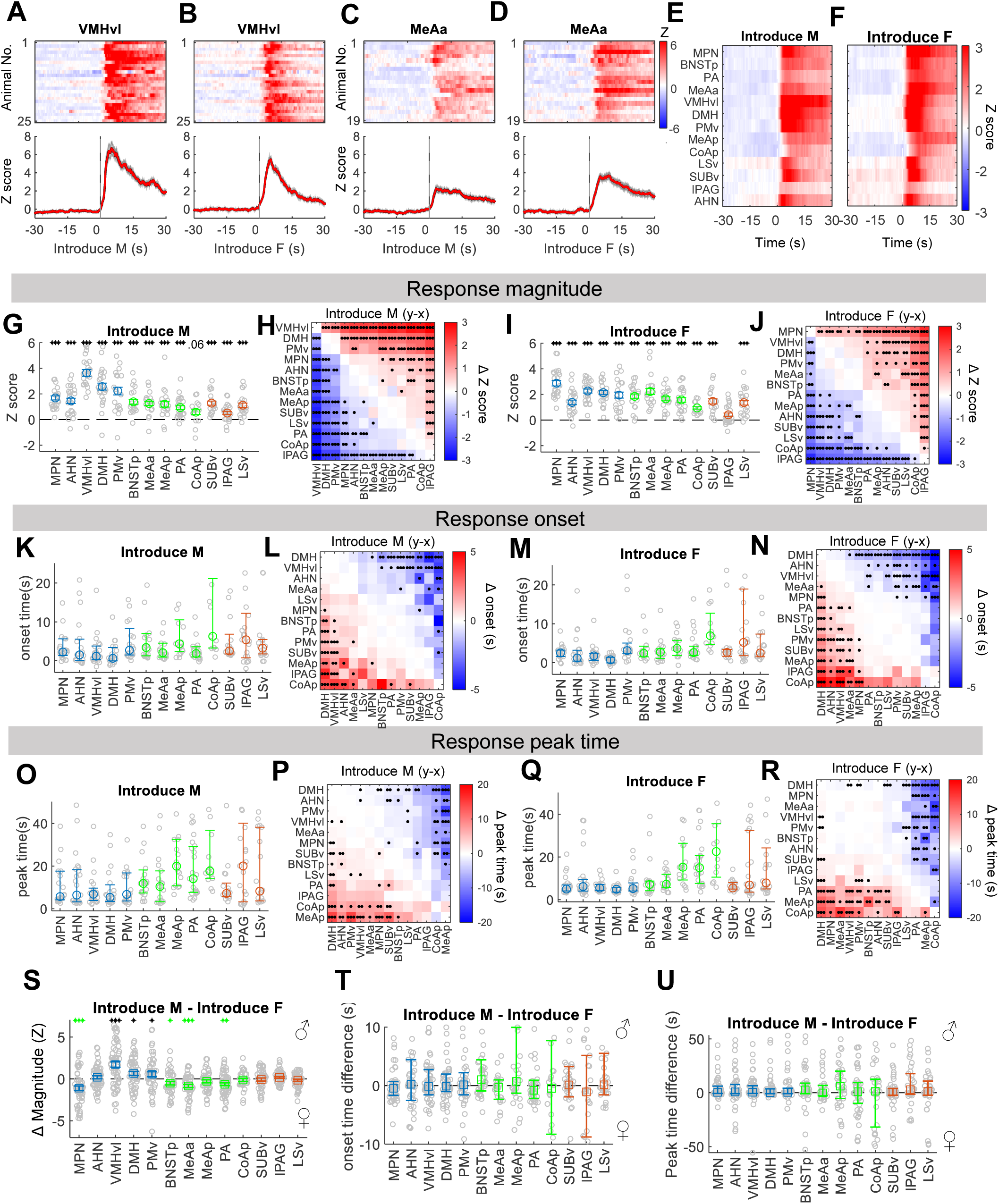
Broad activation of the expanded SBN during initial encounters with male and female intruders. **A-B.** Heatmap (top) and PETHs (bottom) of Z scored ΔF/F signal of the VMHvl aligned to the male **(A)** and female **(B)** intruder introduction from all recording animals. **C-D.** Heatmap (top) and PETHs (bottom) of MeAa Ca^2+^ signals aligned to the male **(C)** and female **(D)** intruder introduction from all recording animals. **E-F.** Heatmaps showing average Z-scored ΔF/F aligned to male **(E)** and female **(F)** introduction across all recorded regions. **G and I.** Average Z-scored ΔF/F during 0-30s after male **(G)** and female **(I)** introduction. n =13-25 animals. **H and J.** Heatmap showing the difference in average Z-scored ΔF/F during male **(H)** and female **(J)** introduction between each pair of regions. n = 21-61 sessions. **K and M.** Average onset of responses upon male **(K)** and female **(M)** introduction. n =12-25 animals. **L and N.** Heatmap showing the difference in average response onset upon male **(L)** and female **(N)** introduction between each pair of regions. n = 17-59 sessions. **O and Q.** Average latency to the peak response after male **(O)** and female **(Q)** introduction. n =13-25 animals. **P and R.** Heatmap showing the difference in average response peak time after male **(P)** and female **(R)** introduction between each pair of regions. n = 17-59 sessions. **S, T, and U**. Differences in response magnitude (Z scored ΔF/F) (**S**), onset time (**T**), and peak time (**U**) during the male and female introduction. n = 28-63 sessions. Shades in **A-D** and error bars in **G, I, and S**: Mean ± SEM; in **M, O, Q, T and U**: Median ± 25%. Each gray circle in **G, I, K, M, O, and Q** represents one animal. Each gray circle in **S-U** represents one recording session. **G, I, S T and U:** one sample t-test (if pass Lilliefors normality test) or Wilcoxon signed-rank test (if not pass Lilliefors normality test) and p values are adjusted with Benjamini Hochberg procedure for controlling the false discovery rate. **H, J, L, N, P, and R:** paired t-test (if pass Lilliefors normality test) or Wilcoxon signed-rank test (if not pass Lilliefors normality test) and p values are adjusted with Benjamini Hochberg procedure for controlling the false discovery rate. *p<0.05; **p<0.01; ***p<0.001. See Table S1 for raw data and detailed statistics.

The temporal dynamics of the responses differed across regions but, interestingly, were similar during male and female entries (**Figure 2K-R**). In general, hypothalamic regions increased activity more rapidly during intruder entry than extra-hypothalamic regions (**Figure 2K-N**). During both male and female introduction, DMH rose with the shortest latency (< 1s), significantly faster than most other regions (**Figure 2K-N**). PMv showed the slowest activity increase among all the hypothalamic regions with a median latency of approximately 3 s (**Figure 2K-N**). Amygdala areas generally took longer to respond to the intruder than hypothalamic areas. MeAa and PA responded in approximately 2-3 s while MeAp took around 4 s to respond after the intruder introduction (**Figure 2K-N**). CoApm responded the most slowly during intruder introduction, with a median response latency over 6 s (**Figure 2K, M**). The activity increase of hypothalamic regions also peaked more rapidly, with average latencies of approximately 5 s, while the responses of amygdala regions took 7-23 s to peak (**Figure 2O-R**). Among the extra-hypothalamic regions, MeAa consistently demonstrated the shortest peak time, faster than MeAp and CoApm. Notably, there was no difference in onset latency or time to peak between male and female intruder introduction, suggesting that, unlike the magnitude of the response, the temporal dynamics of a region’s response are largely independent of the intruder sex (**Figure 2T, U**). Given that the hypothalamic responses during intruder entry are larger and faster than amygdala responses, it is unlikely that the hypothalamic responses result from amygdala inputs, at least initially, despite the fact that all recorded amygdala regions, except CoApm, project directly and densely to the recorded hypothalamic regions (Canteras et al., 1992a, 1995; Dong and Swanson, 2004)

During the introduction of a novel object, AHN, VMHvl, and DMH increased activity slightly but significantly, suggesting that activities in these regions could also be influenced by arousal or novelty (**Figure S3A and B**). However, compared to the conspecific introduction, the response magnitude during object introduction was significantly smaller (**Figure S3C-E**).

### The magnitude but not the sequence of responses differs between male and female investigation

After the initial large Ca^2+^ increase upon intruder introduction, Esr1 cells in all regions in the hypothalamus and amygdala, except AHN, increased activity during investigating males and females (**Figure 3A-K**, S4). During the male investigation, VMHvl showed the largest activity increase, followed by PMv, DMH, and MeAp (**Figure 3C-H**). During the female investigation, MPN, PA and MeAa showed the greatest activity increase, followed by MeAp, BNSTp, VMHvl, and PMv (**Figure 3I-J**). Outside the hypothalamus and amygdala, SUBv showed a slight increase in activity, while lPAG and LSv showed no activity change during either male or female investigation (**Figure 3G, I**). We calculated the difference in response magnitude (Z score) during male and female investigation for each recording session and found that VMHvl, DMH, and PMv (p = 0.06) showed male-biased responses, whereas MPN, PA, and MeAa showed female-biased responses (**Figure 3K**). Although investigating males and females evoked activity increase in the same ten regions, there was no significant correlation between the response magnitude to males and females across regions, suggesting that male and female cues evoked distinct activation patterns in the social network (**Figure 3L**). No activity change existed in any brain region during novel object investigation, suggesting the response is social-specific (**Figure S3F, G**).

**Figure 3:**
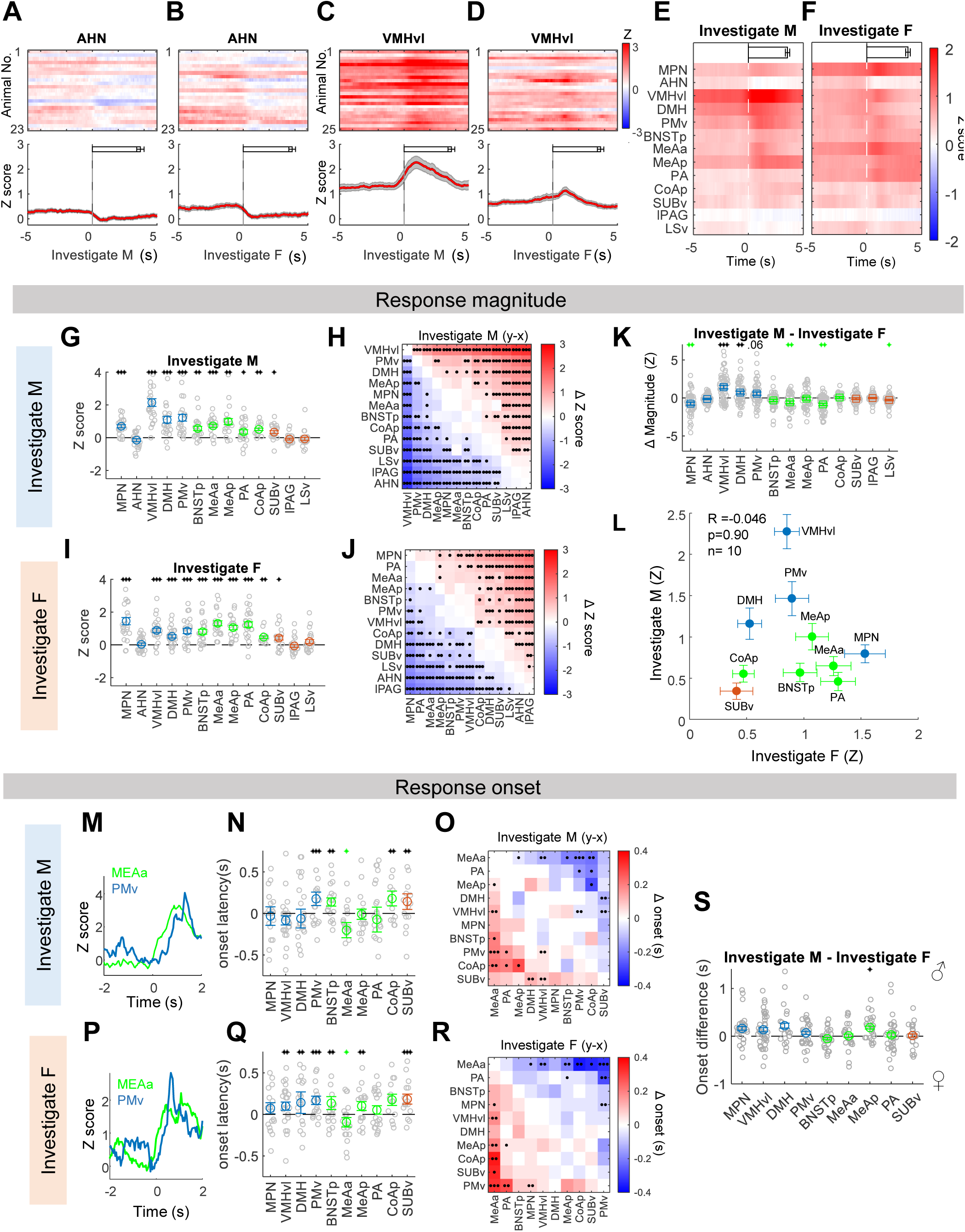
The same regions are activated during male and female investigation but with distinct patterns. **A-B.** Heatmap (top) and PETHs (bottom) of Z scored ΔF/F signal of the AHN aligned to the investigation of male **(A)** and female **(B)** intruders from all recording animals. Horizontal bars indicate investigation duration (mean ± SEM). **C-D.** Heatmap (top) and PETHs (bottom) of VMHvl Ca^2+^ signal aligned to the investigation of male **(C)** and female **(D)** intruders from all recording animals. Horizontal bars indicate investigation duration (mean ± SEM). **E-F.** Heatmaps showing average Z-scored ΔF/F aligned to the investigation of male **(E)** and female **(F)** intruders across all recorded regions. Horizontal bars indicate investigation duration (mean ± SEM). **G and I.** Average Z-scored ΔF/F during the investigation of male **(G)** and female **(I)** intruders. n =13-25 animals. **H and J.** Heatmap showing the difference in average Z-scored ΔF/F during male **(H)** and female **(J)** investigation between each pair of regions. n = 19-60 sessions. **K.** Differences in response magnitude (Z-scored ΔF/F) during the male and female investigation. n = 31-59 sessions. **L.** Scatter plot showing that response magnitude during the male and female investigation is not correlated. n=10 responsive regions. **M and P**. Representative simultaneously recorded Ca2+ traces of MeAa and PMv Esr1 cells during investigating male **(M)** and female **(P)** intruders. **N and Q.** The response latency during male investigation **(N)** and female investigation **(Q)** of all responsive regions. n = 13-25 animals. **O and R.** Heatmap showing the difference in average response onset during male **(O)** and female **(R)** investigation between each pair of regions. n = 32-345 trials. **S.** Differences in response onset during the male and female investigation. n = 21-39 sessions. All error bars and shades in PETHs: Mean ± SEM; Each gray circle in **G, I, N,** and **Q** represents one animal. Each gray circle in **K** and **S** represents one recording session. **G, I, K, N, Q, and S:** one sample t-test (if pass Lilliefors normality test) or Wilcoxon signed-rank test (if not pass Lilliefors normality test) and p values are adjusted with Benjamini Hochberg procedure for controlling the false discovery rate. **H, J, O, and R:** paired t-test (if pass Lilliefors normality test) or Wilcoxon signed-rank test (if not pass Lilliefors normality test) and p values are adjusted with Benjamini Hochberg procedure for controlling the false discovery rate. *p<0.05; **p<0.01; ***p<0.001. Black and green indicate the average values above or below 0, respectively. **L:** Pearson’s cross-correlation. See Table S1 for raw data and detailed statistics.

**Figure 4:**
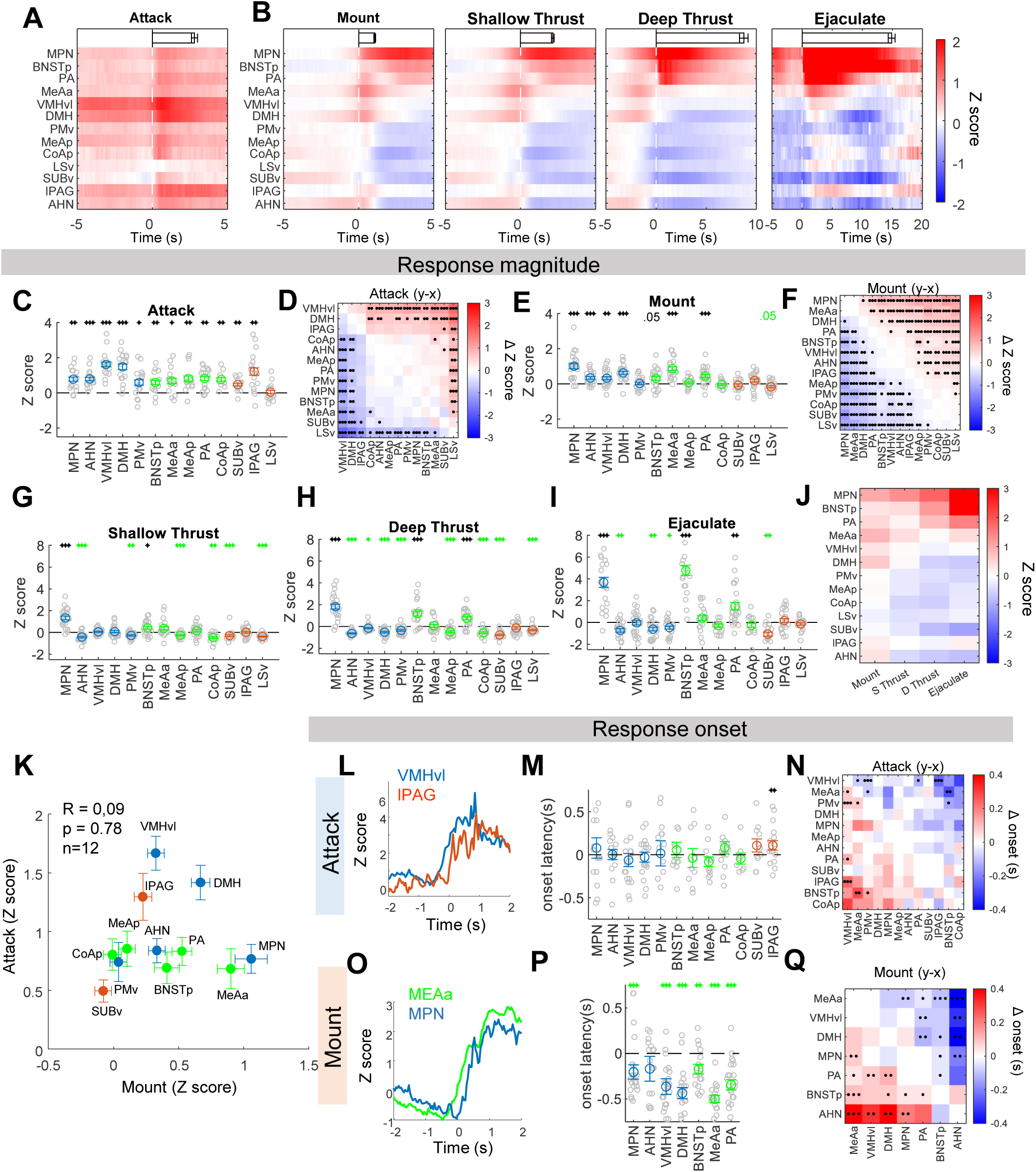
Distinct activation patterns in the expanded SBN during male aggressive and sexual behaviors. **A-B.** Heatmaps showing average Z scored ΔF/F aligned to attack **(A)** and various phases of sexual behaviors **(B)** across all recorded regions. Horizontal bars indicate the average duration of the behavior episodes (mean ± SEM). **C and E.** Average Z scored ΔF/F during attack **(C)** and mount **(E)**. n =8-24 animals. **D and F.** Heatmap showing difference in average Z scored ΔF/F during attack **(D)** and mount **(F)** between each pair of regions. n = 11-55 sessions. **G-I.** Average Z scored ΔF/F during shallow thrust **(G),** deep thrust **(H),** and ejaculation **(I)**. n =12-25 animals. **J.** Heat map showing the average Z scored ΔF/F during various stages of male sexual behaviors across regions. **K.** Scatter plot showing that response magnitude during attack and mount is not correlated. n=12 regions that are responsive during at least one behavior. **L and O.** Representative simultaneously recorded Ca2+ traces of VMHvl and lPAG Esr1 cells during attack (**L**) and MeAa and MPN Esr1 cells during mount (**O**). **M and P.** The response latency during attack **(M)** and mount **(P)** of responsive regions. n = 8-23 animals. **N and Q.** Heatmap showing the difference in average response onset during attack **(N)** and mount **(Q)** between each pair of regions. n = 23-260 trials. All error bars and shades of PETHs: Mean ± SEM; Each gray circle in **C, E, G, H, I, M, and P** represents one animal. **C, E, G-I, M, P:** one sample t-test (if pass Lilliefors normality test) or Wilcoxon signed-rank test (if not pass Lilliefors normality test) and p values are adjusted with Benjamini Hochberg procedure for controlling the false discovery rate. **D, F, N, and Q:** paired t-test (if pass Lilliefors normality test) or Wilcoxon signed-rank test (if not pass Lilliefors normality test) and p values are adjusted with Benjamini Hochberg procedure for controlling the false discovery rate. *p<0.05; **p<0.01; ***p<0.001. Black and green indicate the average values above or below 0, respectively. **I:** Pearson’s cross-correlation. See Table S1 for raw data and detailed statistics.

The temporal response patterns during male and female investigation differed across regions but were largely sex-independent (**Figure 3M-S**). We focused our analysis on the subset of regions that showed a significant increase during the male or female investigation and compared the response onsets of two simultaneously recorded regions during trials when they both increased their activity significantly (Z>2) (**Figure 3M-S**). Unlike the response during intruder entry, MeAa, instead of DMH, was the fastest responding region during both male and female investigation (**Figure 3M-S**). In fact, MeAa was the only region that started to rise slightly before the onset of close investigation, suggesting that its activity change likely does not require direct physical contact (**Figure 3N, Q**). MeAa responded significantly earlier than MPN (p=0.08 for male investigation), VMHvl, and PMv, making it a possible driving force for hypothalamic activation (**Figure 3O, R**). During both male and female investigations, PMv represents one of the slowest regions to respond, although its activity increase was one of the highest, especially during male investigation (**Figure 3G, N, O, Q and** R). Indeed, the fastest responsive region, MeAa, showed only a modest increase in activity during investigating males, suggesting that the response onset and magnitude are largely independent variables (**Figure 3G**). When comparing the response onset during the male and female investigation, only MeAp showed a slightly faster response during male investigation (**Figure 3S**). Thus, the temporal sequence of the responses during social investigation in the expanded SBN is largely invariant to the intruder’s sex and is likely a stable network property.

### Distinct response patterns during attack and sexual behaviors

After a period of investigation, animals expressed distinct actions towards male and female intruders. Specifically, they attacked male intruders and mounted female intruders. Mount refers to the process in which a male tries to grasp the female’s flank with its front paws and establishes an on-top position. Although both attack and mount involve quick movements towards the intruder, the activity pattern in the expanded SBN is highly distinct (**Figure 4**, S4). During attack, the activity increase is widespread, with LSv as the only exception (**Figure 4A, C**). The lack of activity increase in LSv is interesting, given its role in suppressing aggression (Leroy et al., 2018; Wong et al., 2016). VMHvl, DMH, and lPAG were the regions with the highest activity increase, while the remaining responsive regions showed a similarly moderate activity increase (**Figure 4A, C, and D**). It is worth noting that lPAG and AHN only increased activity during attack but not male investigation, suggesting their action-specific responses (**Figure 3G and 4C**). In comparison, the activity increase during mounting was more limited and graded. MPN and MeAa showed the highest activity increase, followed by DMH, PA, BNSTp, VMHvl, and AHN (**Figure 4E, F**). The remaining six regions did not show consistent activity increase (**Figure 4E, F**). The activity change during attack and mounting was not significantly correlated, suggesting that the activation pattern is behavior-specific (**Figure 4K**).

The temporal dynamics of the responses during attack and mount were also distinct (**Figure 4L-Q**). During attack, VMHvl responded the most quickly, increasing activity significantly earlier than MeAa, PA, PMv, and lPAG (**Figure 4L-N**). lPAG was the only region that increased activity after the onset of attack, consistent with its main role in driving biting during attack (**Figure 4M**) (Falkner et al., 2020). But overall, there was relatively little temporal difference in the response onset among the 12 regions that significantly increased activity during attack, suggesting that attack initiation may involve simultaneous activation of many regions in the limbic system (**Figure 4N**).

The activity increase in the hypothalamus and amygdala often preceded the mount onset, possibly partly because mount often follows close interaction with the female (**Figure 4O-P**). Across regions, MeAa responded with the shortest latency, significantly earlier than MPN and PA, while MPN and PA increased activity earlier than BNSTp (**Figure 4P-Q**). Thus, mount appears to involve sequential recruitment of regions in the expanded SBN.

After the male establishes an on-top position, it rapidly (22-25 Hz) and shallowly thrusts with its pelvis (Morali et al., 2003). If the male detects the female’s vagina, mount advances to intromission, or deep thrust, a motion that enables the male to insert its penis into the female’s vagina (Morali et al., 2003). After multiple intromissions, typically 5-20 times, the male ejaculates, characterized by ceased movement and a slow dismount (Hull and Dominguez, 2007). As male sexual behavior advanced, the activities of different regions diverged (**Figure 4G-J, S4**). MPN, BNSTp, and PA gradually increased activity from shallow thrust to ejaculation (**Figure 4G-J, S4**). The activity increase in BNSTp during ejaculation was particularly striking, reaching a peak value 3-4 times higher than during any other period (**Figure 4I, S4**). In contrast, most other regions gradually decreased activity as the animals advanced from mount to deep thrust (**Figure 4G-J, S4**). Towards the end of the deep thrust, there was a widespread suppression of activity in the expanded SBN except for MPN, BNSTp, and PA (**Figure 4H-J, S4**). During ejaculation, some suppressed regions, including VMHvl, MeAa and CoApm and lPAG, showed a slight increase in activity while others, such as SUBv, AHN and DMH, were further suppressed (**Figure 4J, S4**). Altogether, these results revealed distinct activation patterns during male aggression and sexual behaviors: the former is characterized by a widespread and simultaneous activation across many regions, while the latter features robust activation of a small set of regions and a gradual suppression of many others.

### Delineation of the aggression-biased network and the mating-biased network

We then performed the principal component analysis (PCA) based on the mean responses (Z score) of all regions during various phases of social behaviors (**Figure 5A**). The variance in responses across behaviors could be explained nearly fully (99%) by the first four principal components (PCs) (**Figure 5B**). PC1 is composed of responses during male sexual behaviors, especially ejaculation and deep thrust. MPN and BNSTp have the highest PC1 score, followed by PA and MeAa, while all other regions have negative PC1 scores, consistent with their suppressed activity during advanced sexual behaviors (**Figure 5C, D**). PC2 is dominated by responses during male-directed behaviors, especially male investigation, and VMHvl is the region with the highest score, followed by DMH, PMv, and MeAp (**Figure 5C, D**). PC3 is again composed of female-directed behavior, but unlike PC1, mount and shallow thrust have the highest loadings, while ejaculation has a negative loading (**Figure 5C**). MeAa and MPN are the regions with the highest PC3 scores, followed by PA, consistent with their activity increase during mount (**Figure 5D**). Finally, PC4 features a high loading during attack and negative loadings during investigation (**Figure 5C**). lPAG shows the highest PC4 score, followed by DMH, VMHvl, and AHN (**Figure 5D**).

**Figure 5.**
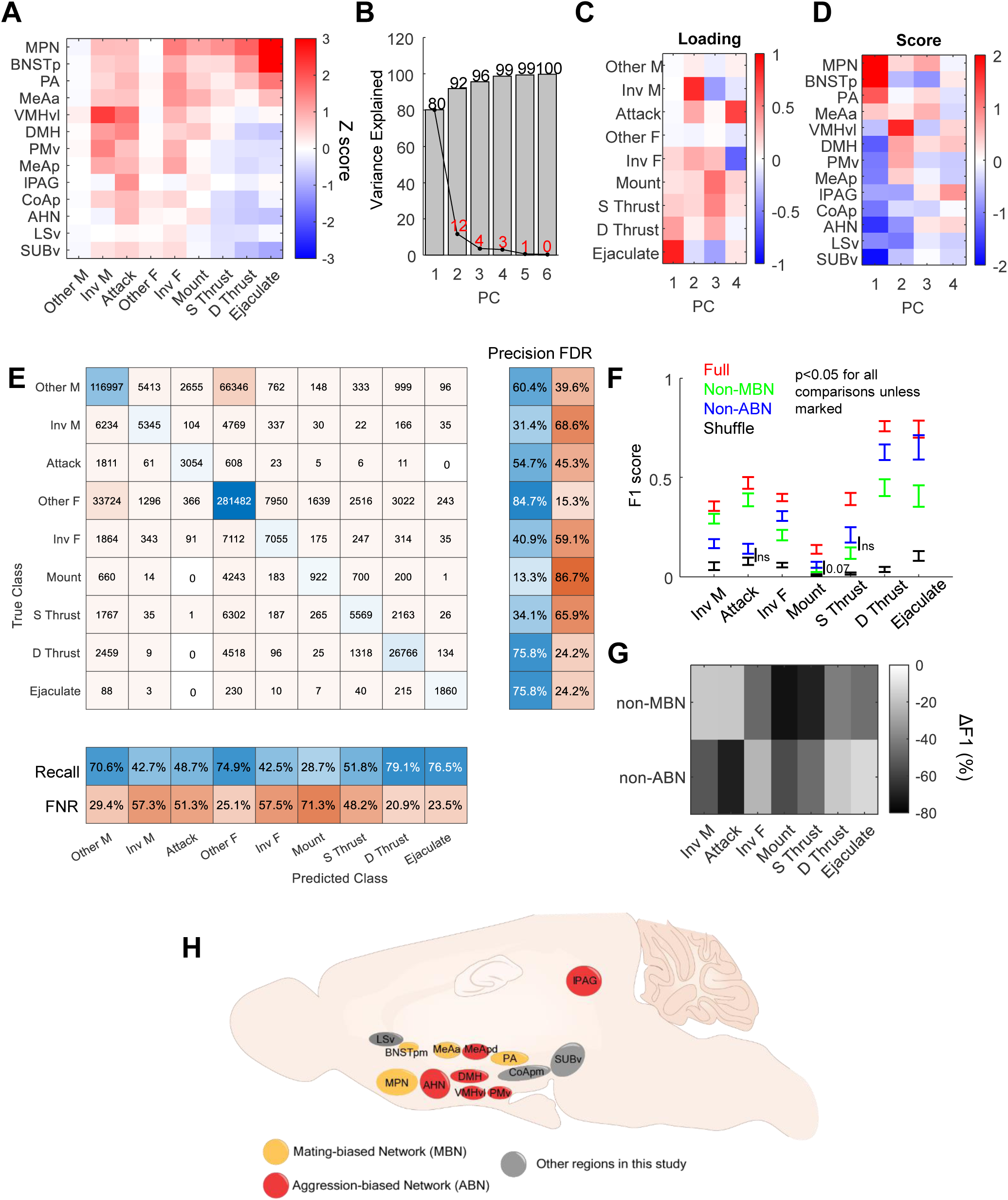
Activities in MBN and ABN predict male sexual and aggressive behaviors, respectively. **A.** Heat map showing the average Z scored ΔF/F during male- and female-directed social behaviors across regions in male mice. “Other M” and “Other F” refer to periods when the male or female intruder is present but no specific social behavior is annotated. Inv: investigate; S Thrust: shallow thrust; D Thrust: deep thrust. **B.** The variance in responses during different behaviors explained by the first 6 PCs. **C.** The loading (coefficient) of the first 4 PCs. **D.** The scores of the first 4 PCs for each region. **E.** Confusion matrix shows the number of frames that are correctly and incorrectly classified for each behavior across all sessions. Left columns show the precision (blue) and false discovery rate (FDR, orange). Bottom rows show the recall (blue) and false negative rate (FNR, orange). **F.** F1 scores for various behaviors computed using full models that includes data from all recording regions, non-MBN models, non-ABN models, and models built with shuffled data. Error bar: mean ± SEM. Paired t-test (if pass Lilliefors normality test) or Wilcoxon signed-rank test (if not pass Lilliefors normality test). All comparisons with p values < 0.05 unless marked. ns: not significant. n = 17-24 animals. **G.** Heat map showing the averaged decrease in F1 score when using non-MBN and non-ABN models compared to the full model. See Table S1 for raw data and detailed statistics.

Based on this analysis, we propose an aggression-biased network (ABN) and a mating-biased network (MBN) among our recorded regions (**Figure 5H**). ABN contains six brain regions, including VMHvl, PMv, MeAp, DMH, AHN, and lPAG. AHN and lPAG are preferentially activated during attack and thus represent the motor output nodes of ABN. On the other hand, MBN contains four regions, including MPN, BNSTpm, PA, and MeAa. MeAa is mainly activated during the early phase, while BNSTp increases activity preferentially during the late phase of copulation, while MPN and PA are activated throughout sexual behaviors.

To further understand the distinctiveness of the activation pattern during each behavior and the contribution of ABN and MBN activity to predict the behavior, we trained a discriminant analysis model for each recording session using 80% of randomly selected data points (training set). Then we predicted the behavior categories of the remaining 20% of data points (testing set). When the model was trained and tested using only frames that were annotated with a specific behavior (16% of total frames across all videos), it successfully separated the behaviors based on the neural activation patterns and predicted most behaviors accurately (F1 score (mean across behaviors ± STD) = 0.81 ± 0.11) (Figure S5). Mount and shallow thrust have relatively low F1 scores as they are sometimes misclassified as each other, possibly due to their close temporal proximity and the difficulty for humans to determine the precise transition point (**Figure S5B, C**).

We next trained and tested the model using all frames, including frames without specific social behaviors (annotated as “other”) (**Figure 5E-G**). These “other” frames mainly involve two animals being far apart but also contain instances when animals were close but showed no discrete actions. After including these unspecified frames, we found that F1 scores (mean ± STD = 0.49 ± 0.22) were decreased for all social behavior categories, although the all-frame model was able to predict all behaviors significantly better than models trained with shuffled recording data (**Figure 5E, F and S5C**). The decrease in F1 score was mainly driven by misclassifying “other” frames as ones with specific behaviors and *vice versa* (**Figure 5E**). Interestingly, the drop in the F1 score varied widely across behaviors. For deep thrust and ejaculation, F1 scores only dropped slightly (<15%), while the F1 score for mount dropped by nearly 70% (**Figures 5F** and **S5C**). This result suggests that neural activation patterns associated with deep thrust and ejaculation are highly distinct and rarely occur outside of these behaviors during social interaction. In contrast, the activity pattern associated with mount is much less so, possibly reflecting many intended mounts that are not manifested behaviorally.

We then investigated the contribution of ABN and MBN activity in predicting the behaviors by training models using only data from non-MBN regions or non-ABN regions. When using the non-MBN model, F1 scores for all female-directed behaviors, but not male-directed behaviors, dropped dramatically compared to the full model trained using data from all regions, supporting a key role of MBN activity in determining the behavior output towards the females (**Figure 5F, G**). When using the non-ABN model, F1 scores for male-directed behaviors dropped the most, but notably, F1 scores for mount and shallow thrust also decreased, suggesting that mount initiating could require activity in both MBN and ABN (**Figure 5F, G**). It is worth noting that the activity of non-MBN and non-ABN regions could still predict male- and female-directed behaviors better than shuffled controls, suggesting that although ABN and MBN are preferentially involved in aggression and mating, respectively, they are not exclusive for the behavior (**Figure 5F**).

### Strengthened functional connectivity in the expanded SBN during the initiation of social behaviors

To address whether the functional connectivity among regions in the expanded SBN changes with social behaviors, we next calculated the coefficient of determination (R^2^) using 1-s moving windows (25 data points) between each pair of simultaneously recorded regions (**Figure 6A**). The time window was chosen based on the typical duration of a behavior episode (approximately 1-15s, see **Figure 3A-F and 4A-B**). Varying time windows from 0.4 s to 2 s did not change the result qualitatively. To remove the auto-correlation between adjacent data points, we pre-processed the data by calculating the 1^st^ order derivative of each recording trace as the difference between adjacent data points (Figure S6). Figure 6 shows the correlation between PMv and VMHvl as an example. For the representative recording session, PMv-VMHvl had R^2^ of approximately 0.09 at the baseline before intruder introduction (**Figure 6B, D**). In the presence of a male intruder, the R^2^ jumped to approximately 0.15, further increased to 0.25 during male investigation, and peaked at around 0.44 during attack (**Figure 6B-D**). PMv-VMHvl correlation also increased in the presence of a female intruder and was further elevated during female investigation (**Figure 6D**). However, the correlation gradually decreased as the sexual behaviors advanced and reached a level below the pre-intruder baseline during deep thrust and ejaculation (**Figure 6C, D**). These changes are consistent across recording sessions from different animals: the functional connectivity, i.e., R^2^, between PMv and VMHvl strengthened during male-male interaction and peaked during attack, whereas it gradually decreased during sexual behaviors (**Figure 6E-M**).

**Figure 6.**
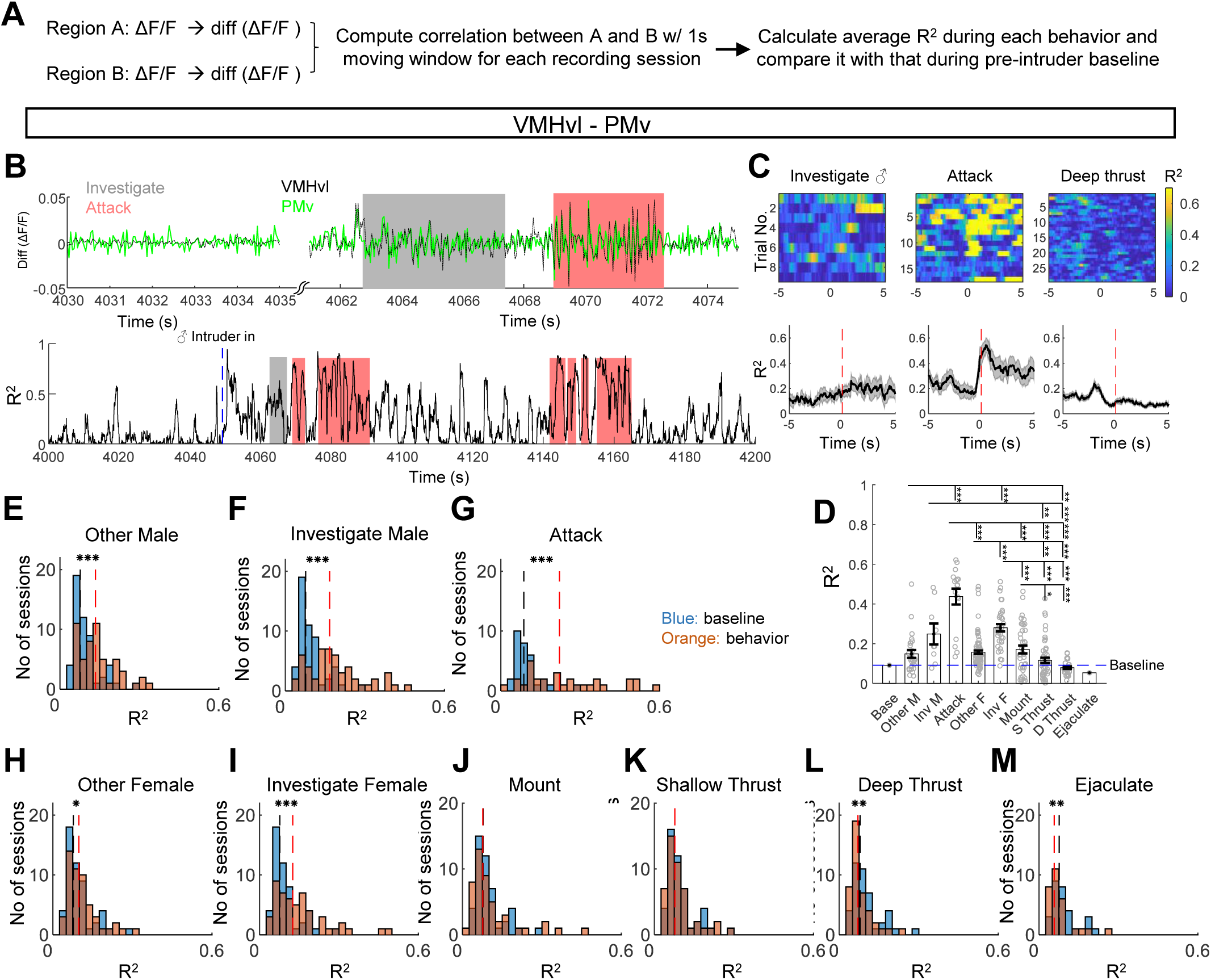
Changes in functional connectivity between VMHvl and PMv during various social behaviors. **A.** The procedure to calculate the coefficient of determination (R^2^) between a pair of regions during various behaviors. **B.** Differential Z scored ΔF/F traces of VMHvl (black) and PMv (green) (top) and their moment-to-moment R^2^ (bottom) from a representative recording session. Shades indicate behavior episodes. **C.** Heatmaps (top) and PETHs (bottom) aligned to the onset of male investigation (left), attack (middle), and deep thrust (right). It is from the same recording session, as shown in B. **D.** Average R^2^ between VMHvl and PMv during various behaviors in the recording session shown in **B** and **C**. n = 1-86 trials. “Base” refers to pre-intruder period. “Other M” and “Other F” refer to periods when the male or female intruder is present but no specific social behavior is annotated. Inv: investigate; S Thrust: shallow thrust; D Thrust: deep thrust. Kruskal-Wallis test followed by Benjamini Hochberg procedure for controlling the false discovery rate. **E-M**. Histograms show the distribution of R^2^ at the pre-intruder baseline (orange) and during specific behavior epochs (blue). Black and red dashed lines indicate the median values of the R^2^ during the baseline and behavior periods. Baseline and behavior sessions are matched. Paired t-test (if pass Lilliefors normality test) or Wilcoxon signed-rank test (if not pass Lilliefors normality test). Error bars and shades of PETHs: Mean ± SEM. *p<0.05; **p<0.01; ***p<0.001. See Table S1 for raw data and detailed statistics.

One crucial question is whether the correlation change during social behavior simply reflects changes in motor output. To understand the relationship between functional connectivity and movement, we calculated R^2^ for all pairs during the low (bottom 25%) and high-velocity (top 25%) periods when the animal was alone in the cage. Although no pair of regions showed a significant change in R^2^ with velocity, we noticed that connections involving lPAG, SUBv, BNSTpm, PA, and LSv tended to increase strength during the high-velocity period (**Figure S7A-C**). To further address the question, we identified the time points when the locomotion initiated (reach peak speed >8 pixel/fr) after a quiescence period (mean speed < 1 pixel/fr for > 1s) and constructed PSTHs of R^2^ aligned to the movement onset (**Figure S7D, E**). This analysis confirmed that functional connectivity between regions involving lPAG, SUBv, BNSTpm, PA, and LSv increased after movement initiation. However, functional connections among hypothalamic regions, medial amygdala, and cortical amygdala were largely invariant to movement (**Figure S7F and G**). Thus, although movement may contribute to changes in functional connectivity between some regions, it is not expected to play a significant role in modulating the functional connectivity among the hypothalamus, medial and cortical amygdala, e.g., between VMHvl and PMv, during social behaviors.

We then examined the functional connectivity across all 78 pairs of regions during different social behaviors and reached several general conclusions. First, some regions showed a significant correlation in activity even at the baseline level (**Figure 7A and B**). This baseline correlation may be considered resting-state connectivity. The strongest functional connectivity was observed among amygdala regions, including MeApd, PA, CoApm and SUBv, or connections involving lPAG (**Figure 7A and B**). After male and female introduction but in the absence of social interaction, the functional connectivity pattern remained largely the same except for strengthening of the VMHvl and PMv connection, especially in the presence of a male intruder (**Figure 7A**, C **and F**). During the social investigation with males, the correlation among VMHvl, PMv, and DMH, increased significantly (**Figure 7A, D**). During the female investigation, there is a broader, weak increase in functional connectivity (**Figure 7A, G**). The most striking changes in correlation occur during attack and mount, the social behavior initiation phase. During attack, the functional connectivity between 95% of pairs in the social network increased significantly and drastically (**Figure 7A, E, and L-N**). Intriguingly, the increase in functional connectivity does not necessarily require a net change in Ca^2+^ activity. For example, although LSv did not show an increase in response during attack, its functional connectivity with the rest of the expanded SBN nevertheless increased (**Figure 7A, L**). During mounting, there is also an overall increase in connectivity in the network, but interestingly not for connections involving the VMHvl (**Figure 7A, H and L-N**). As the sexual behavior advanced, we observed a general decorrelation in the network (**Figure 7A, I-N**). During shallow thrust, the connectivity largely returned to the baseline level except for several weakly strengthened connections involving BNSTpm, MPN, PA, lPAG, and AHN (**Figure 7A, I, L**). During deep thrust and ejaculation, most connections weakened, some reaching a level significantly below the pre-intruder baseline (**Figure 7A, I-N**).

**Figure 7.**
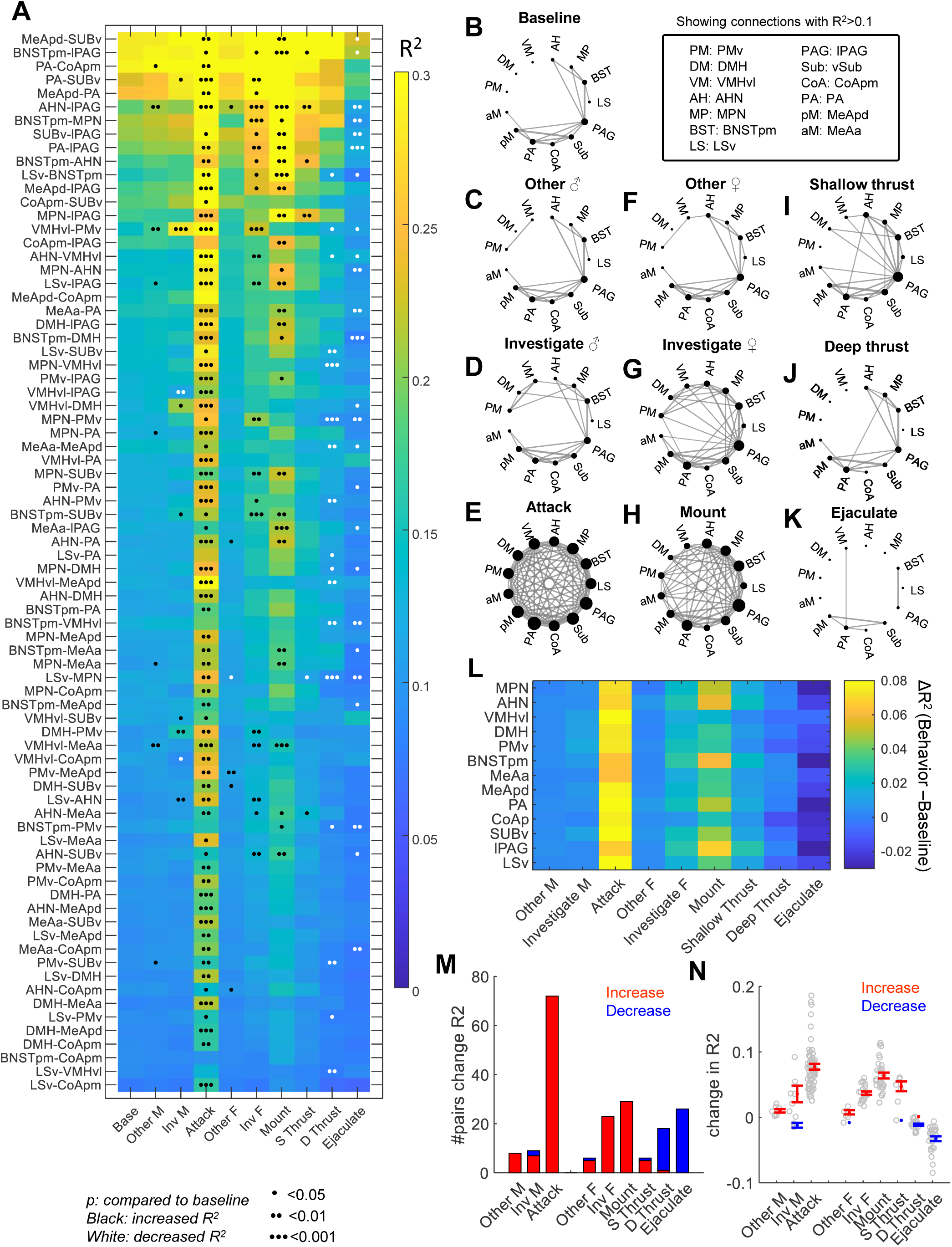
Changes in functional connectivity across the expanded SBN during social behaviors. **A.** Heat map shows the average R^2^ of all pairs of regions during various behavior epochs and its comparison to the R^2^ values during the baseline period. “Base” refers to pre-intruder period. “Other M” and “Other F” refer to periods when the male or female intruder is present but no specific social behavior is annotated. Inv: investigate; S Thrust: shallow thrust; D Thrust: deep thrust. Paired t-test (if pass Lilliefors normality test) or Wilcoxon signed-rank test (if not pass Lilliefors normality test). P values are adjusted using with Benjamini Hochberg procedure for controlling the false discovery rate. *p<0.05; **p<0.01; ***p<0.001. Black and white indicate a significant increase or decrease from the baseline, respectively. **B-K.** Graph plots showing the strength of functional connectivity (R^2^) among different regions during various social behavior epochs. Only connections with R^2^> 0.1 are shown. The size of a node reflects its overall connection strength. **L.** Heat map shows averaged change of R^2^ of each region with all other regions during various behaviors. **M.** the number of pairs of regions that show significantly increased R^2^ (red) or decreased R^2^ (blue) from the pre-intruder baseline. **N.** Change in R2 values from the pre-intruder baseline for significantly changed connections. Red and blue show the mean ± SEM of significantly increased and decreased connections during each behavior. n = 0-72 pairs of regions. See Table S1 for raw data and detailed statistics.

Introducing a small jitter (40-200 ms) into the simultaneously recorded Ca^2+^ traces abolished the across-region correlation during all behaviors, suggesting that the correlation is highly sensitive to the precise alignment of activities between regions (**Figure S8**). Altogether, these results suggest that initiating social actions involves coordinated activation across the social network, including regions that appear to show no net change in activity. Intriguingly, advanced sexual behaviors are accompanied by a “dissociated” brain state.

## Discussion

Mating and fighting are two complicated behaviors supported by the coordinated activation of many brain regions. Here, using a modified multi-site optical recording system, we examined the neural activities across multiple regions in the limbic system during sexual and aggressive behaviors in male mice. These recordings revealed widespread activities in the network that evolved in distinct ways over the course of male-male and male-female social interactions.

### The overall activation pattern in the expanded SBN during male social behaviors

Here, we characterized three aspects of the Ca^2+^ responses during social behaviors: magnitude, timing, and functional connectivity among regions (**Figure S9**). Overall, the response magnitude was behavior- and intruder sex-specific, whereas the timing was behavior- but not intruder sex-specific. Functional connectivity in the network increases drastically during the fast action phase of social behaviors, i.e., attack and mount, and decreases as the males engage in advanced copulation.

The most prominent responses of most recorded regions occurred when the animal first encountered the intruder. This is perhaps not surprising as the net change of sensory cues was the largest at that moment. An unexpected finding is that the entry response was generally higher and faster in the hypothalamus than amygdala, possibly reflecting a higher level of information convergence and a lower spontaneous activity in the hypothalamus. DMH was consistently the first (<1s) to respond regardless of the intruder sex, and this response occurred before the physical interaction of the two animals (median latency of first interaction: 3.7 s for a male intruder and 3.6 s for a female intruder), suggesting its potential role in the initial detection of distant social targets. Several regions in the ABN, such as VMHvl, DMH, and PMv, showed higher responses to males than females, while all MBN regions, including MeAa, PA, BNSTpm, and MPN, showed the opposite preference. Thus, information regarding the sex identity of the intruder is represented widely in the expanded SBN quickly after the intruder’s presence.

Ten out of thirteen regions were activated significantly during social investigation, all responding to both males and females. The response magnitude in each region showed a sex bias similar to that during initial intruder encounters. Interestingly, MeAa instead of DMH was the first region to increase its activity during each episode of social investigation, and it was also the only region to respond before the investigation onset. Like MeAp, MeAa receives extensive inputs from the accessory olfactory bulb and projects densely to the medial hypothalamus and other olfactory-related amygdala regions (Pardo-Bellver et al., 2012). However, it has received much less attention regarding its functions in social behaviors, as immediate early gene studies found that MeAa activation is not social behavior-specific. Handling, tail pinch, mating, and fighting activate the area similarly (Kollack-Walker and Newman, 1995), leading to the hypothesis that MeAa belongs to a general arousal circuit (Newman et al., 1997). Here, we found the MeA^Esr1^ cells were rapidly and specifically activated during social investigation, especially towards females, but not object investigation, suggesting a potential social-specific role. Interestingly, in comparison to MeAp, MeAa projects more densely to the ventral striatum and ventral tegmental area, regions essential for moment-to-moment social interest (Dai et al., 2022; Gunaydin et al., 2014; Pardo-Bellver et al., 2012).

The activity change during attack and mount differed vastly. Attack was accompanied by activity increases in all regions except the LSv, a region that has been found to suppress aggression. VMHvl demonstrated the fastest and largest response during attack. In particular, its activity increase preceded PA, PMv, and PAG, regions that have been shown to play roles in aggression, suggesting that VMHvl may be the “ignitor” of attack (Chen et al., 2020; Falkner et al., 2020; Stagkourakis et al., 2020; Stagkourakis et al., 2018; Yamaguchi et al., 2020; Zha et al., 2020). However, VMHvl does not mediate attacks on its own. The functional connectivity of nearly the entire expanded SBN strengthened drastically during attack. Thus, although the attack signal may originate from the VMHvl, its sustenance likely requires the entire network. As a result, modulating many nodes in the network could cause changes in attack (Chang and Gean, 2019; Chen et al., 2020; Falkner et al., 2020; Hong et al., 2014; Leroy et al., 2018; Stagkourakis et al., 2018; Unger et al., 2015; Wong et al., 2016; Yamaguchi et al., 2020; Zelikowsky et al., 2018; Zha et al., 2020).

Mount was accompanied by a relatively limited activity increase in the network, involving seven activated regions vs. twelve during attack. MPN and MeAa demonstrated the highest activity increase. Interestingly, the activity rise in the MeAa is earlier than MPN, the most-established region for male sexual behavior. While it is possible that MeAa activity increase partially reflects changes related to social investigation given that social investigation often precedes mount, we did not find MeAa to respond the most quickly during attack, which also often follows close interaction. The functional importance of MeAa in sexual behaviors remains to be examined. At the network level, the functional connectivity between many regions also increased during mounting, as in the case of attack, but connections involving VMHvl remained largely unchanged. This perhaps explains a lack of deficits in mounting when VMHvl was artificially inhibited, even though VMHvl is one of the regions with increased activity during mounting (Lee et al., 2014; Lin et al., 2011). As sexual behavior advances, we saw a clear divergence of activity patterns across regions. PA, MPN, and BNSTp were the only regions that gradually increased their activity, peaking during ejaculation. All other regions gradually decreased their activity, although some increased activity slightly during ejaculation. The neural activity pattern during advanced sexual behavior was highly distinct and could be used to predict the behaviors reliably. During advanced sexual behaviors, the most striking change at the network level was an overall decrease in functional connectivity. No connection, regardless of whether it is between regions responsive or not, is strengthened, and many decrease below the baseline level. This result suggests that the brain enters a “dissociated” state during advanced mating. Interestingly, during copulation, male rats are reported to have drastically reduced sensitivity to pain (Gonzalez-Mariscal et al., 1992; Szechtman et al., 1981). Although the response threshold to other external cues has not been studied systematically, anecdotally, we found that male mice appear oblivious to the external world during deep thrust. Whether these behavior changes result from decreased communication across brain regions remains to be investigated in future studies.

### The mating-biased network

Here, we propose the MBN for supporting male sexual behaviors, which includes MPN, BNSTp, PA, and MEAa. These regions, except MeAa, showed a consistent increase during all stages of male sexual behaviors, with higher responses in more advanced stages. The activity increase was likely caused by a combination of sensory inputs (e.g., olfactory and somatosensory) and internal cues (e.g., hormone and neuromodulator) and may have been used to drive moment-to-moment actions associated with mating.

MPN is unarguably the most studied region for male sexual behaviors. Since 1941, numerous studies have demonstrated that damage in MPN impaired or abolished male sexual behaviors without spontaneous recovery, suggesting its irreplaceable role in mating (Brookhart and Dey, 1941). Both c-Fos and *in vivo* recordings corroborate functional results (Wei et al., 2018). Our recordings confirm the large increase in MPN activity throughout the male sexual behaviors. However, we do notice some heterogeneity in MPN responses across animals. In animals with optic fibers located in posterior MPN, the cell responses to females tended to be weaker than those with anterior MPN-targeted fibers (although all animals were included in the analysis). Our recent study suggests that posterior MPN is mainly activated when facing a social threat and plays a vital role in suppressing aggression against a superior opponent (Wei et al., Accepted).

The role of BNSTpm in male sexual behavior has been long suspected based on the remarkably dense c-Fos expression after male sexual behaviors (Coolen et al., 1997; Kollack-Walker and Newman, 1995; Kollack and Newman, 1992; Newman et al., 1997). However, lesion-induced behavioral deficits are anything but remarkable. Lesioned animals can execute the whole sequence of male sexual behaviors, although the intromission interval and the number of intromissions preceding ejaculation increases (Claro et al., 1995; Emery and Sachs, 1976; Powers et al., 1987; Valcourt and Sachs, 1979) (but Bayless et al. (2019) report poorer sexual behavior performance after inhibiting BNSTpm aromatase cells). The BNSTpm^Esr1^ cell responses may explain the relatively minor behavior deficit. As these cells are mainly activated during ejaculation, the most important function of the cells could be related to ejaculation, such as initiation of ejaculation, ejaculation-induced sexual satiation, or hormone changes.

PA has largely escaped the attention of neuroscientists, possibly due to its relatively low c-Fos induction after sexual behaviors (Kollack-Walker and Newman, 1997). Nevertheless, our recent functional studies demonstrated that PA*^Esr1^* cells are both necessary and sufficient for male sexual behaviors through their projection to the MPN (Yamaguchi et al., 2020). Most strikingly, when the PA^Esr1^ to MPN projecting cells are inhibited, males rarely mount and never achieve deep thrust (Yamaguchi et al., 2020).

The role of MeAa in sexual behaviors remains elusive. Early studies found that MeAa lesions could abolish all aspects of male sexual behaviors (Kondo, 1992). However, the MeApd, but not MeAa, expressed c-Fos specifically after exposure to female pheromone cues and consequently became the focus of many studies (Fernandez-Fewell and Meredith, 1994). However, recent cell type-specific ablation argued against the important role of MeApd in male sexual behaviors (Unger et al., 2015). Consistent with those functional results, we found no significant activity increase of MeAp^Esr1^ cells during male mating. Given the strong and rapid activation of MeAa during the early phase of male sexual behavior, future studies are needed to investigate its function in male sexual behaviors.

### The aggression-biased network

The proposed ABN contains AHN, DMH, VMHvl, PMv, MeAp, and lPAG based on their preferential responses during aggression over sexual behaviors. Here, we will summarize their known functional roles in aggression and highlight the new insights revealed by our recordings.

VMHvl has now been firmly established as a critical site for conspecific aggression (Lee et al., 2014; Lin et al., 2011; Yang et al., 2013). Consistent with its central role in aggression, VMHvl shows the largest response during male introduction, investigation and attack. In particular, VMHvl is the first to respond and one of the top regions to increase functional connectivity during attack, supporting its central role in attack initiation.

PMv is a major input to the VMHvl and is highly responsive to conspecific olfactory cues (Chen et al., 2020; Motta et al., 2013). Indeed, PMv is among the most activated regions during male investigation. However, PMv response during attack is relatively weak and significantly slower than the VMHvl, suggesting its secondary role in initiating attack. It also suggests that the rise of VMHvl activity during attack is unlikely a result of olfactory inputs channeling through the PMv.

DMH has only been studied recently for its role in aggression. Zelikowsky *et. al.* found that tachykinin-expressing DMH cells are both necessary and sufficient for social isolation-induced aggression in male mice, although the neural response of the cells during attack has not been reported (Zelikowsky et al., 2018). Here, we found that DMH, just like VMHvl, showed a male-biased response during all stages of social behaviors. Whether DMH influences aggression mainly through its connection with VMHvl or its parallel projection to the midbrain, e.g., PAG, remains to be investigated.

Several recent studies demonstrated a necessary and sufficient role of MeApd GABAergic cells in inter-male aggression (Hong et al., 2014; Miller et al., 2019; Nordman et al., 2020; Padilla et al., 2016; Unger et al., 2015), although the cell *in vivo* Ca^2+^ responses during attack remain unreported. Consistent with the functional results, we found that MeApd^Esr1^ cells increased their activity during both attack and male investigation, with attack evoking a higher response than mount. MeApd is also a region with substantially increased functional connectivity during attack, especially with VMHvl, supporting its important role in attack initiation.

Consistent with a role in motor action, lPAG^Esr1^ cells showed increased activity exclusively during attack. We previously found that lPAG is a key downstream of VMHvl for attack initiation, although stimulating VMHvl-lPAG pathway only induces attacks with low efficiency (Falkner et al., 2020). Indeed, most regions in the ABN project to lPAG to some extent (Beitz, 1982), and we speculate that lPAG may serve as a common gateway that integrates inputs from ABN to initiate action. Thus, blocking lPAG is sufficient to block attack, whereas activating any specific input only triggers attack weakly or not at all (Falkner et al., 2020).

AHN was considered a part of the aggression circuit mainly based on the functional evidence acquired in hamsters (Ferris et al., 1997; Gobrogge et al., 2007). However, later, AHN was proposed to be a part of the predator defense circuit instead of the social behavior circuit, given its strong c-Fos activation after predator encounters (Canteras, 2002; Martinez et al., 2008). Consistent with this hypothesis, Xie *et. al.* recently reported that AHN GABAergic cells (the main population) bi-directionally control defensive attack against predators (e.g., snakes), but have little influence on conspecific aggression in mice (Xie et al., 2020). Here, we found that AHN^Esr1^ cells uniquely increased activity during the action phase of social behaviors, especially attack, calling for further investigation of AHN’s role in fighting and mating.

Although PA is considered a part of the mating circuit, it is important to note that PA does also play a role in male aggression through its projection to VMHvl (Stagkourakis et al., 2020; Yamaguchi et al., 2020; Zha et al., 2020). PA indeed shows a consistent increase during both male investigation and attack, although the responses during female investigation and sexual behaviors are significantly higher. Both aggression- and sexual behavior-relevant cells in PA express *Esr1* but are largely distinct at the single-cell level (Yamaguchi et al., 2020).

LSv is unique in that it is the only region that did not increase activity during attack. It is also the only region whose connection with VMHvl is not significantly strengthened during attack. LS has been recognized as a region that “gates” aggression for decades. Early lesion studies demonstrated that LS damage could cause “septal rage”, i.e., unprovoked ferocious attack (Albert and Chew, 1980). More recent studies confirmed that LS could negatively modulate aggression at least partly through its GABAergic projection to VMHvl (Leroy et al., 2018; Wong et al., 2016). In light of these functional results, the lack of activity increase in LSv may signal “permission” to attack.

In summary, we investigated Ca^2+^ responses in the expanded SBN during social behaviors in male mice. Our results suggest that the sex identity information is broadly represented across the expanded SBN. Fighting and mating are associated with highly distinct patterns of activation in the network. Attack features synchronized activation of many regions in the limbic system with VMHvl being the potential ignitor. Sexual behavior is associated with sequential activation of a small set of regions and their gradual changes during behavior progression. The network activity during advanced copulation is particularly unique, featuring strong activation of three regions and suppression of others, and reduced communication across regions. These results provide a holistic view regarding the neural generation of social behaviors and will serve as an important guide for future functional studies.

## STAR methods

### Experimental model and subject details

#### Animals

Experimental mice for MFP recording were socially naïve, Esr1-2A-Cre male mice (10–24 weeks, Jackson stock no. 017911). After surgery, all test animals were single-housed. Intruders used were group-housed BALB/c males or group-housed C57BL/6 females (both 10–36 weeks, Charles River). Mice were housed at 18–23 °C with 40–60% humidity and maintained on a reversed 12-h light/dark cycle (dark cycle starts at 10 a.m.) with food and water available ad libitum. All experiments were performed in the dark cycle of the animals. All procedures were approved by the IACUC of NYULMC in compliance with the NIH guidelines for the care and use of laboratory animals.

## Method details

### Optical setup

The optical setup was a modified version of a typical fiber photometry setup (Falkner et al., 2016) according to a previously described FIP system (Kim et al., 2016). Briefly, blue LED light (Thorlabs, M470F1, LEDD1B) was bandpass filtered (Semrock, FF02-472/30-25), reflected on a dichroic filter (Semrock, FF495-Di03-25×36), and coupled into a custom-designed 19-fiber multi-fiber bundle (Doric Lenses, BFP(19)_100_110_1100-0.37_4m_FCM-19X) through a 10x objective (Olympus PLN). Emission light was bandpass filtered (Semrock, FF01-535/50) and projected onto the CCD sensor of a camera (Basler, acA640-120um) via an achromatic doublet (Thorlabs, AC254-060-A-ML). The connector end of the fiber bundle was imaged by the camera. The LED was driven by DC current, and the optical power out of the tip of every single fiber was set to be ∼30 μW. The sampling rate of the camera was 25 frames per second.

### Stereotaxic surgery

Esr1-2A-Cre mice were anesthetized with 1.5%-2% isoflurane and placed on a stereotaxic surgery platform (Kopf Instruments, Model 1900). 60-100 nl AAV2-CAG-FLEX-GCaMP6f viruses (Vigene, custom prepared) or AAV1-CAG-FLEX-GCaMP6f viruses (Addgene, 100835-AAV1, 3x dilution) were delivered unilaterally into each of the following targeted brain regions as described previously (Fang et al., 2018): LSv (AP 0.05, ML -0.65, DV -3.25); MPN (AP 0.00, ML -0.33, DV -4.90); BNSTpm (AP -0.30, ML -0.80, DV -3.60); AHN (AP -1.05, ML -0.55, DV -5.15); DMH (AP -1.75, ML -0.55, DV -5.2); VMHvl (AP -1.70, ML -0.75, DV -5.80); PMv (AP -2.40, ML -0.55, DV -5.70); MeAa (AP -1.10, ML 2.10, DV -4.90); MeAp (AP -1.60, ML 2.10, DV -4.92); PA (AP -2.35, ML 2.20, DV -4.92); CoApm (AP -2.85, ML 2.90, DV -5.20); SUBv (AP -3.35, ML 2.60, DV -4.60); lPAG (AP -4.90, ML -0.45, DV -2.40). Each regional data included in the final analyses had correct virus expression and fiber tip position as verified by histology.

The multi-fiber arrays were constructed using MT Ferrules (US Conec, No 12599) and 100 µm-core optic fibers (Doric Lens, NA0.37) as described previously (Sych et al., 2019). For each animal, two custom-made multi-fiber arrays were implanted, one designed to target seven medial regions on the left side and the other to target five lateral regions on the right side. In the same animal, a custom-made optic-fiber assembly targeting lPAG (Thorlabs, CFX126-10) was also implanted. All optic fibers are targeted ∼250 μm above the injection sites and secured using dental cement (C&B Metab ond, S380). lPAG fiber was implanted at (AP -5.20, ML -0.45, DV -2.00) after tilting the head 8 degrees down rostrally to avoid collision with the other two arrays. Lastly, a 3D-printed plastic ring for head fixation was cemented on the skull (Osborne and Dudman, 2014).

### Behavioral analysis and tracking

Behaviors were recorded under dim room light via two cameras from top and side views (Basler, acA640-100gm) using StreamPix 5 (Norpix), which also coordinated the MFP camera in synchrony. Behaviors were then manually annotated and animal positions were tracked on a frame-by-frame basis using custom software in MATLAB (https://pdollar.github.io/toolbox/). Annotated behaviors are defined as follows: ‘Investigation’, the resident mouse made nose contact with either the facial or anogenital region of the intruder mouse or the whole body of the toy mouse; ‘Attack’, a suite of actions initiated by the resident toward the male intruder, which included lunges, bites, tumbling and fast locomotion episodes between such behaviors; ‘Mount’, began when the resident male charged toward the rear end of the female body, rose and grasped the female’s flank with his forelimb, and ended by aligning his body with the female’s and assuming the on-top posture; ‘Shallow thrust’, the male grasped the female’s body tightly with his forelegs and made rapid shallow pelvic thrusts; ‘Deep thrust’, deep rhythmic movement of pelvis presumably with penile insertion into the vagina; ‘Ejaculation’, the male froze at the end of an intromission event while continuously clutching onto the female and then slumping to the side of the female. Ejaculation occurred only once in a female session, signaling the end of sexual behaviors. It was always confirmed by the presence of a vaginal plug after the recording. Behavioral annotations were made by trained experimenters, during which neural responses were not available to the experimenter. The animals were tracked using custom MATLAB software (https://github.com/pdollar/toolbox)(Burgos-Artizzu et al., 2012; Lin et al., 2011). The velocity of the animal was calculated as the distance between the animal’s body center locations in adjacent frames (pixels/s).

### Multi-fiber photometry recording

The recording started three weeks after the virus injection. For each recording session, the head-mounted MT ferrules were connected to the matching connectors at the end of the optic fiber bundle (Doric lens, BFP(19)_100/110/1100-0.37_4m_SMA-19x). A drop of liquid composite (Henry Schein, 7262597) was applied to the outer part of the junction and cured with blue LED curing light (Amazon) to stabilize the connection. The baseline signal was checked in the absence of the intruder for at least two days to ensure that the signal reached a stable level (<10% difference across days for all regions). On the recording day, after 10 minutes of the baseline period, a sexually receptive female mouse was introduced until the recording male achieved ejaculation or after 60 minutes. Then, 5 minutes after removing the female, a group-housed non-aggressive Balb/C was introduced for 10 minutes. For some recording sessions, 5 minutes after removing the male, a novel object (15 mL plastic tube) was introduced for 10 minutes. Each animal was recorded 2-4 times, with at least three days in between. The order of male and female presentations was counterbalanced across sessions.

### Data analysis

Regions of interest (ROIs) for selected channels were drawn on the grayscale image of the optic fiber bundle, and the average pixel intensity for each ROI was calculated as a readout of the raw Ca^2+^ signal (Fr^aw^) for the region. We then used the MATLAB function ‘‘msbackadj’’ with a moving window of 10% get the flatted signal F_flat_. Then the instantaneous baseline signal was obtained as “F_baseline_ = F_raw_ – F_flat_”. The ΔF/F was then calculated as “ΔF/F = (F_raw_ –F_baseline_)/F_baseline_”. The ΔF/F signal was then Z-scored using the entire recording trace for each channel. All analyses were based on Z-scored ΔF/F.

The response magnitude of each behavior for each recording animal was calculated by first averaging Z-scored ΔF/F during all episodes of the behavior in a session and then averaging the values across all recording sessions of an animal. The difference in response magnitude between two regions was calculated as the difference of average responses of two simultaneous recorded regions of a session and analyzed across all sessions. To compare responses during male-directed and female-directed behaviors (e.g., male investigation vs. female investigation), we calculated the magnitude difference between the behavior towards the male intruder and the female intruder of each session and analyzed all sessions. To calculate the onset of the response of behavior, e.g., attack, we first selected responsive trials when the Z-scored ΔF/F reached >2. For those trials, we then determined the trough time preceding the peak response after smoothing the trace with a low pass filter (threshold 4 Hz). If a session contained at least three responsive trials (Z>2), the average onset time of the behavior of the session was computed. Otherwise, the onset would be registered as NaN. For comparing the onset time during male-directed and female-directed behavior, e.g., male investigation vs. female investigation, we determined the average onset during each behavior in one session and calculated their difference. To compare the onset time of two different regions in one behavior, we identified trials where both regions showed a peak response >2, determined response onset for each, and calculated the difference. We then performed a two-sided t-test (if data passed the normality test) or Wilcoxon signed-rank test (if data did not pass the normality test) on the onset time difference using all trials with a null hypothesis that the onset difference was 0. The peak time during introduction was determined as the latency to reach the maximum value in the first 30s after intruder introduction. The peak time difference between the two regions was calculated based on the peak time of simultaneously recorded traces. The PETHs were constructed by aligning the Z-scored ΔF/F to the onset of a behavior, averaging across trials, averaging across sessions for each animal, and then averaging across animals.

Principal Components Analysis (PCA) was performed using the MATLAB function “pca.” The data submitted to PCA was a 13 x 9 matrix (corresponding to 13 regions and 9 behaviors) whose *i*^th^, *j*^th^ element is the response magnitude (Z scored ΔF/F) of the *i*^th^ region during the *j*^th^ behavior, averaged first over trials, then sessions, and finally subjects. The first four components explained over 99% of the variance.

A linear discriminant model for each recording session was constructed using the MATLAB function “fitcdiscr ”using 80% of randomly selected data (training data). The model was then used to predict the behaviors associated with the remaining 20% (testing data) of the Ca^2+^ recording data in each session using MATLAB function “predict”. We used either all the frames (Figure 5) or the frames annotated with specific social behaviors (Figure S5) for training and testing. For reach session, only channels with correct targeting were used for training and testing the model. The confusion matrix was constructed based on all the testing data from all sessions of all animals. F1 score was calculated as (2 × precision × recall) / (precision + recall) for each behavior and each recording session and averaged across sessions. To calculate the F1 score of shuffled data, the recording traces of all channels were shifted by a random offset (0 to the duration of the recording session) and used for constructing the discriminant model and predicting the behaviors. This procedure was performed once for each session.

For assessing the contribution of MBN and ABN regions to behavior prediction, we constructed the discriminant models based on recordings from non-MBN regions or non-ABN regions and used the model to predict the behaviors. For Figure 5F and Figure S5C, F1 scores were computed separately for each subject by concatenating results across sessions.

To determine the instantaneous coefficient of determination (R^2^) between two regions, we first calculated the first derivative of the Z-scored ΔF/F trace as the difference between adjacent data points (25 points/sec). We then computed the moving-window correlation (window size: 25 data points) using the MATLAB function “movcorr” and its elementwise squaring as R^2^. We then calculated the average R^2^ during each behavior for each recording session and the average of all sessions. To determine whether R^2^ changed significantly during a behavior, we performed paired t-test (if data passed the normality test) or sign test (if data did not pass the normality test) between the averaged R^2^ during the behavior and that during the baseline period of the same session across all sessions. The p values were adjusted using Benjamini & Hochberg procedure for controlling the false discovery rate (FDR). The graph plot for each behavior was generated using MATLAB function “graph”. The averaged R^2^ values during the behavior for all pairs of regions (76 in total) were used as the weights of connections. Only connections with R^2^ >0.1 were shown. The size of the node is proportional to the accumulated weight (R^2^) of the connections involving the node. To determine the importance of the temporal alignment on the correlation between regions, we added a slight jitter (randomly selected from 40, 80, 120, 160 and 200 ms) to one of the Z-scored ΔF/F traces in each pair of regions and then calculated the R^2^ of all pairs.

To determine the relationship between the movement velocity and correlation between regions, we tracked the animal, and calculated its body center velocity and the average R^2^ in the frames with the top 25% movement velocity and those with the bottom 25% velocity for each session. We then calculated the difference between each session’s low-velocity and high-velocity periods and the average across sessions. To determine the onset of movement during the baseline period, we determined troughs and peaks in the velocity trace and selected troughs that precede peaks reaching at least 8 pixels/fr and follow >1s of quiescence (mean velocity < 1pixel/fr). We then constructed PSTHs of R^2^ of each pair of regions aligned to the movement onset in each session, and calculated the average for each session and the average PSTHs of all sessions. We then calculated the difference in averaged R^2^ between post- (0 -1 s) and pre-movement (-1 - 0 s) based on the PSTHs. The movement-sensitive pairs (red-filled circles in Figure S7F and red trace in Figure S7G) are pairs with a ΔR^2^> 0.01.

### Histology and imaging

Mice were over-anesthetized with isoflurane and transcardially perfused with cold 1x phosphate buffered saline (PBS) followed by cold 4% paraformaldehyde (PFA) in 1x PBS. Heads with implants were post-fixed in 4% PFA for at least 72 h at 4°C and then transferred into 15% sucrose solution for 48 h, after which brains were carefully extracted and put into 15% sucrose solution at 4°C overnight. Brains were embedded in OCT mounting medium, frozen on dry ice, and cut into 50 μm-thick sections using a cryostat (Leica). Sections were collected in a 6-well plate, washed three times with 1x PBS, and counter-stained with DAPI (1:20,000; Thermo Fisher, D1306) diluted in PBS-T (0.3% Triton X-100 in 1x PBS) for 15 min. After washing with PBS-T once, sections were mounted on Superfrost slides (Fisher Scientific, 12-550-15) and cover-slipped for imaging via a slide scanner (Olympus, VS120). 10x fluorescent images were acquired to access fiber placements and virus expressions.

### Quantification and statistical analysis

All statistical analyses were performed using MATLAB 2021a (MathWorks) or Prism 9 (GraphPad). All datasets were tested for normality with the Lilliefors test, whenever applicable. Parametric tests, including one-sample t-test, paired t-test, and ordinary one-way ANOVA and multiple-comparison post hoc tests were used if distributions passed the normality test. If distributions failed the normality test, non-parametric tests, including one-sample Wilcoxon signed rank test, Wilcoxon signed rank test, and Kruskal-Wallis test, were used. P values for all multiple one-sample t-tests, one-sample Wilcoxon signed rank tests, multiple pairs of t-tests, and post-hoc multiple-comparison tests were adjusted using Benjamini & Hochberg procedure for controlling the false discovery rate. Significance in all statistical results was indicated as follows: * p < 0.05, **p < 0.01, and ***p < 0.001. Error bars were presented as mean ± s.e.m if most datasets in a figure plot passed the normality test. Otherwise, error bars were presented as median ± 25%. No statistical methods were used to predetermine sample sizes, but our sample sizes were similar to or larger than those reported previously. Statistical details and sample size can be found in the figure legends and Table S1.

