## Supplementary Figures for "Neural dynamics in the limbic system during male social behaviors"

### Figure S1

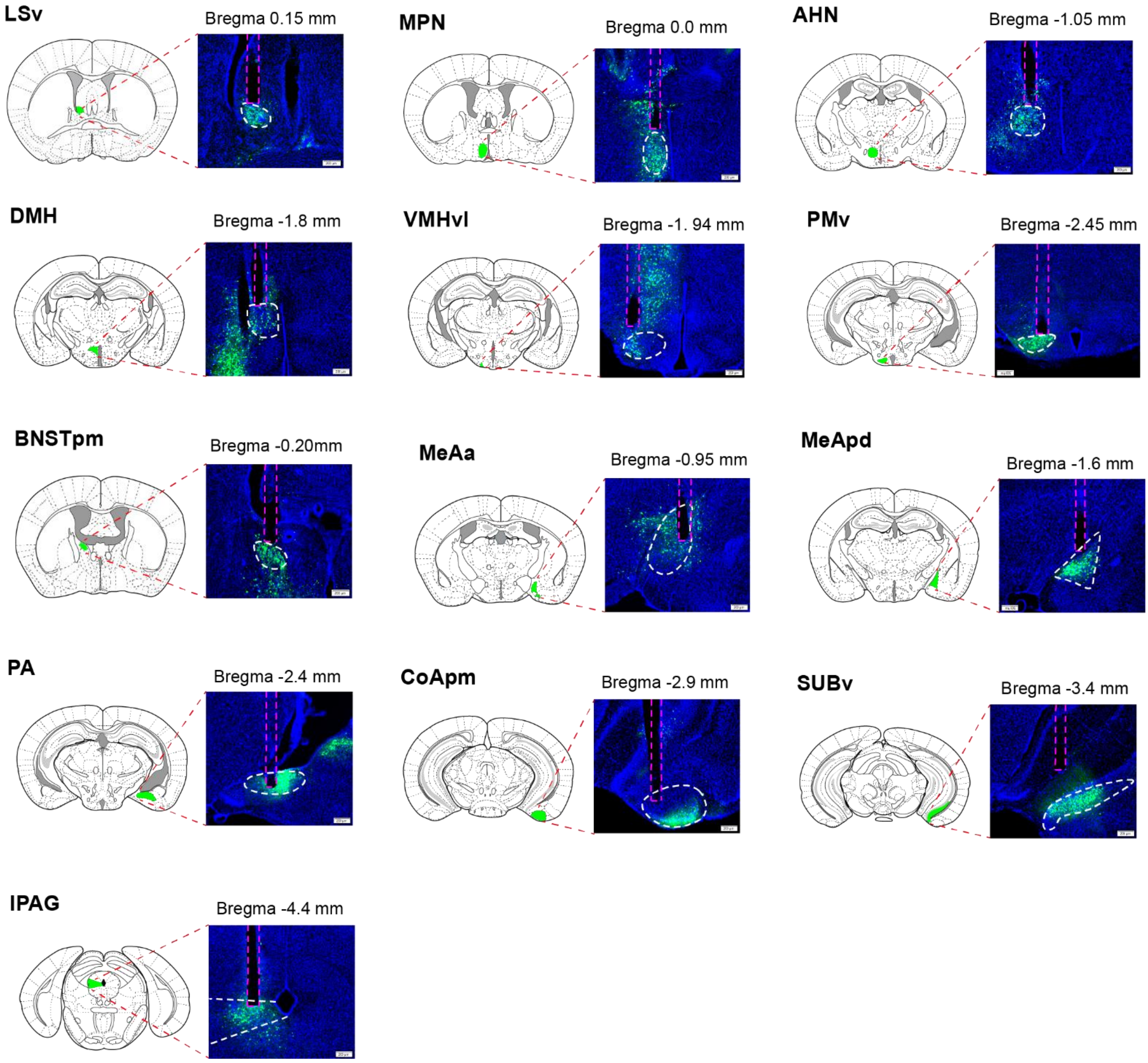


**Figure S1. Representative GCaMP6f expression and optic fiber tracks (red dashed lines) in the 13 targeted brain regions. Related to Figure 1**. Scale bars: 200 µm. The coronal plane at the appropriate Bregma level of the mouse brain atlas (Paxinos and Franklin, 2008) is shown on the left.

### Figure S2

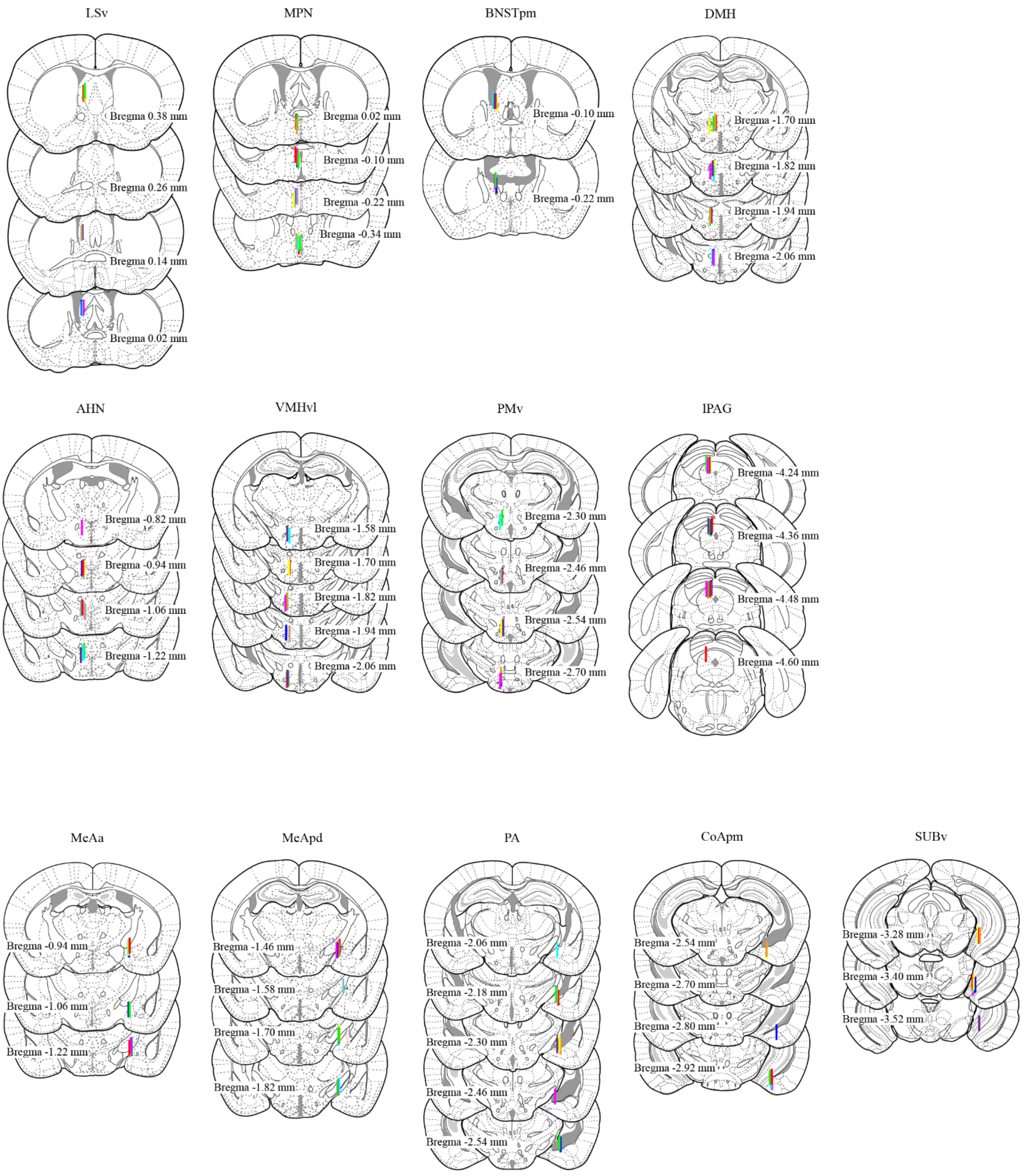


**Figure S2. Summary of the optic fiber locations at each brain region for all recorded animals. Related to Figure 1.** The coronal planes are adopted from (Paxinos and Franklin, 2008).

### Figure S3

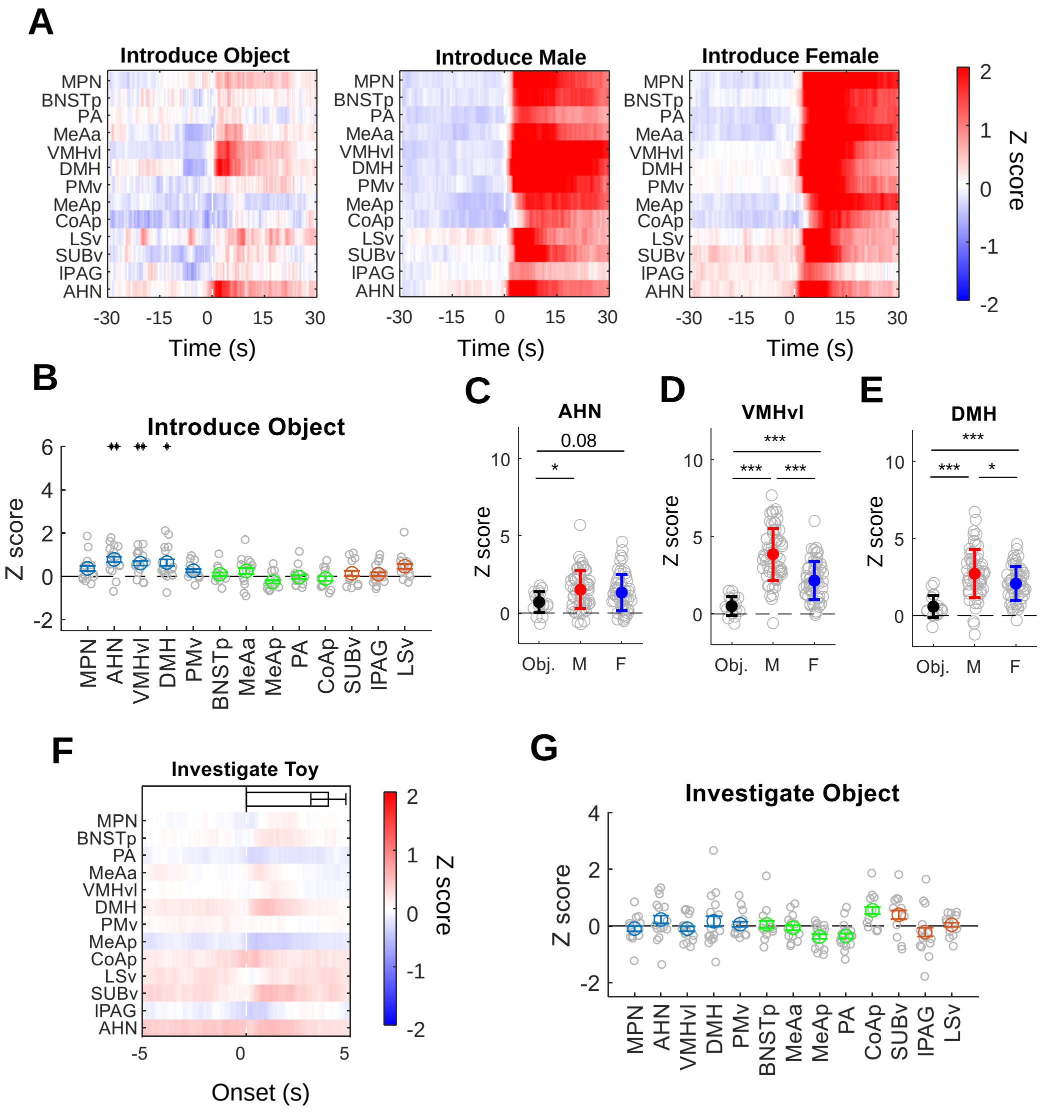


**Figure S3. Changes in Ca2+ activity of regions in the expanded SBN during novel object introduction and investigation. Related to Figures 2 and 3.**

**A.** Heat maps showing average Z scored ΔF/F of various recording regions aligned to the introduction of an object (left), male mouse (middle), and female mouse (right).

**B.** Average Z scored ΔF/F during the first 30 s after object introduction. n =10-15 animals.

**C.** Comparison of averaged Z scored ΔF/F of $AHN^{Esr1}$ cells during object, male and female introductions. n = 18-58 sessions.

**D.** Comparison of averaged Z scored ΔF/F of $VMHvl^{Esr1}$ cells during object, male and female introductions. n = 18-64 sessions.

**E.** Comparison of averaged Z scored ΔF/F of $DMH^{Esr1}$ cells during object, male and female introductions. n = 18-59 sessions.

**F.** Heat maps showing average Z scored ΔF/F of various recording regions aligned to object investigation. Horizontal bars indicate investigation duration (mean ± SEM).

**G.** Average Z scored ΔF/F during object investigation. n =10-15 animals.

All error bars: Mean ± SEM; Each gray circle in **B and G** represents one animal. Each gray circle in **C-E** represents one recording session.

**B and G:** one sample t-test (if pass Lilliefors normality test) or Wilcoxon signed-rank test (if not pass Lilliefors normality test). **C-D:** Ordinary one-way ANOVA followed by multiple comparison tests. E: Kruskal-Wallis test followed by multiple comparison tests. All p values are adjusted with Benjamini Hochberg procedure for controlling the false discovery rate. *p<0.05; **p< 0.01; ***p<0.001. See Table S1 for raw data and detailed statistics.

### Figure S4

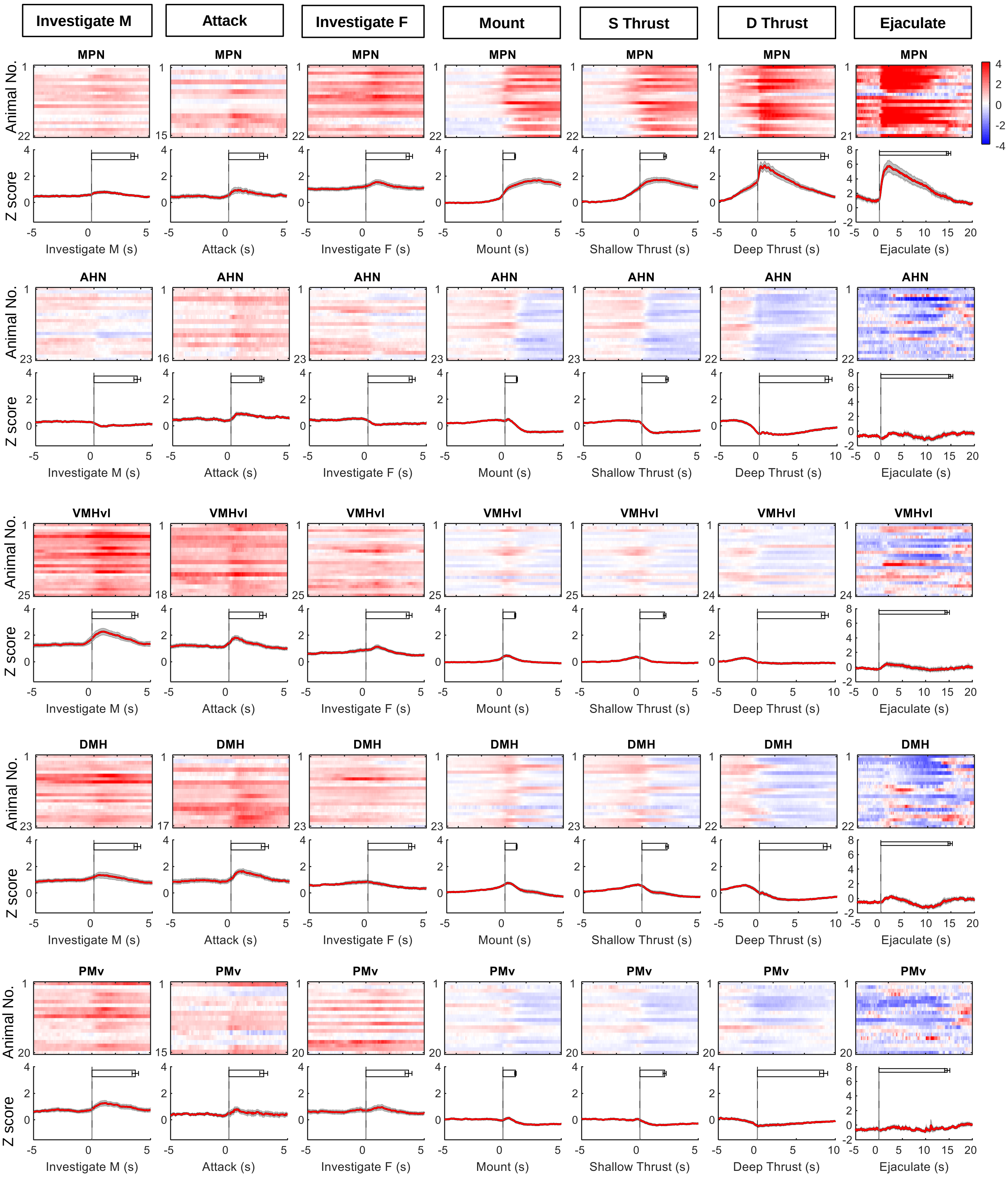

Investigate M
Attack
Investigate F
Mount
S Thrust
D Thrust
Ejaculate
MPN
AHN
VMHvl
DMH
PMv
Animal No.
Z score
Investigate M (s)
Attack (s)
Investigate F (s)
Mount (s)
Shallow Thrust (s)
Deep Thrust (s)
Ejaculate (s)

### Figure S4- continue

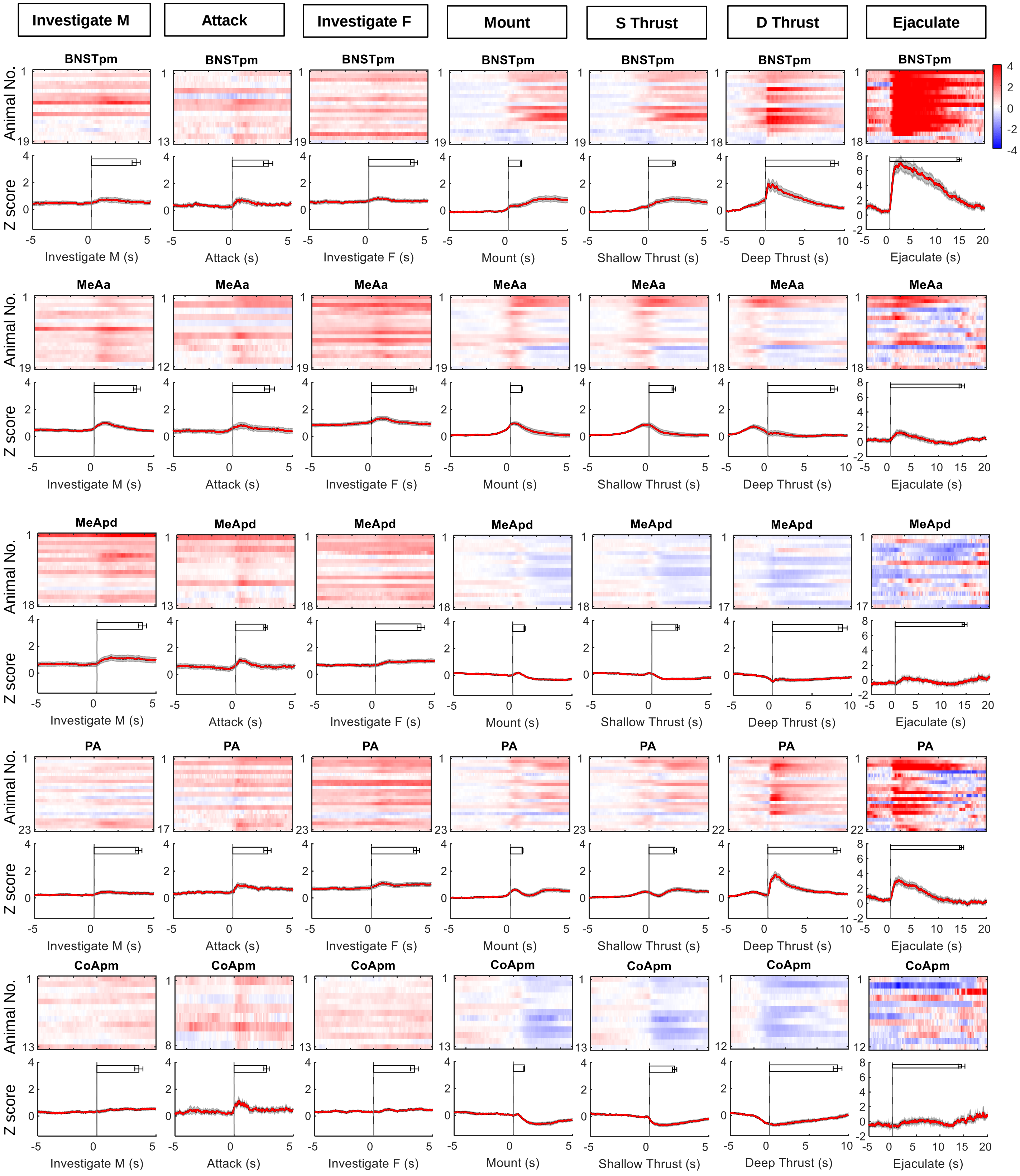

### Figure S4 -continue

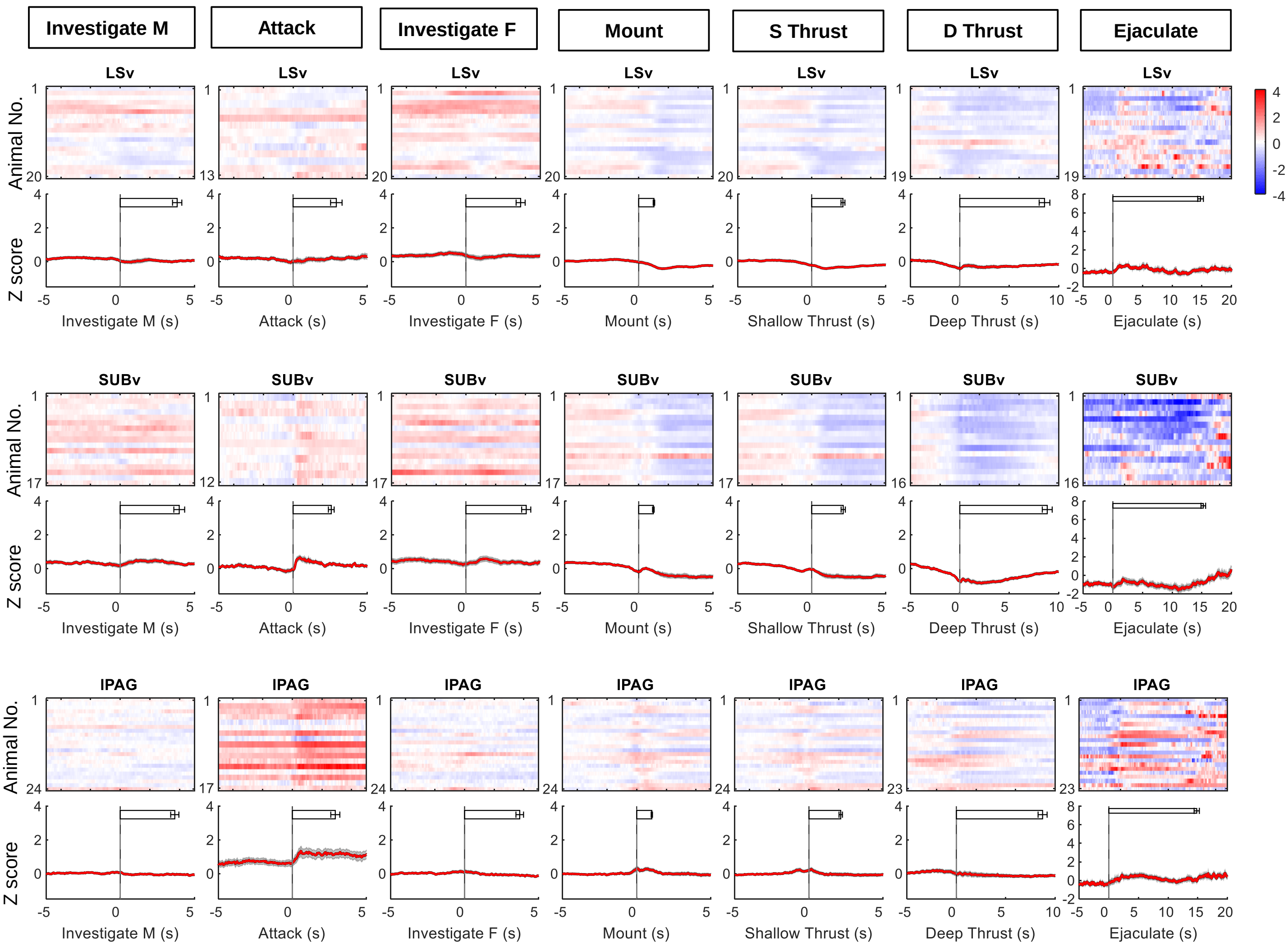


**Figure S4. Heat maps and PETHs of Z scored ΔF/F $Ca^{2+}$ signal aligned to the onset of various behaviors for all recorded regions across all animals. Related to Figures 2, 3, and 4.** The color scale for all heatmaps is the same. Note that the PETH y-axis for ejaculation is different from other behaviors for better visualization. Horizontal bars indicate the average duration of the behavior (mean ± SEM). Shades of PETHs represent ± SEM.

### Figure S5

A

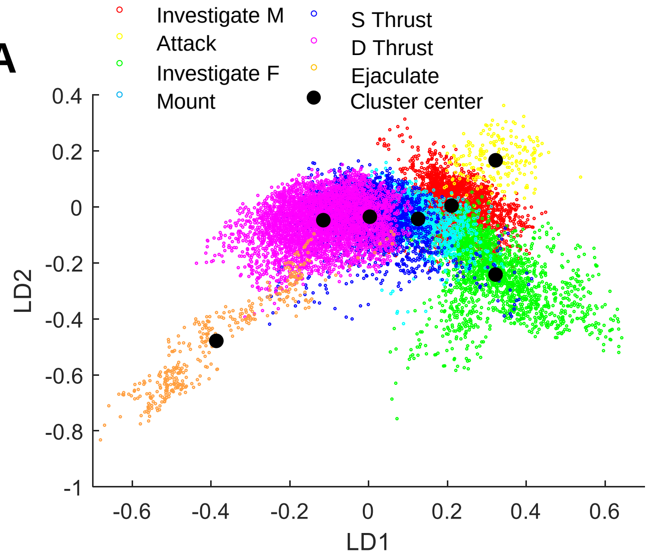


C

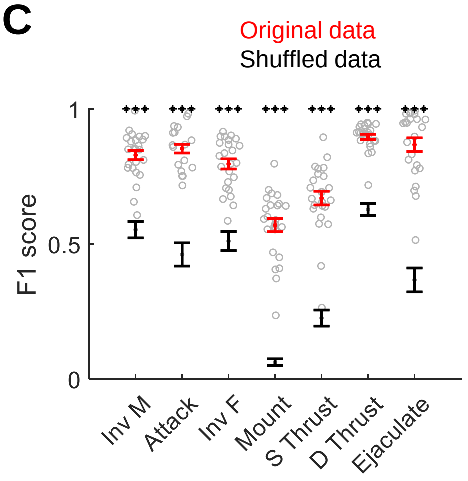


B

| True Class \ Predicted Class | Investigate M | Attack | Investigate F | Mount | Shallow Thrust | Deep Thrust | Ejaculate | Precision | FDR |
|---|---|---|---|---|---|---|---|---|---|
| Investigate M | 13973 | 359 | 1163 | 375 | 416 | 661 | 43 | 82.2% | 17.8% |
| Attack | 281 | 4956 | 188 | 66 | 96 | 30 | 0 | 88.2% | 11.8% |
| Investigate F | 1144 | 148 | 14122 | 512 | 691 | 678 | 41 | 81.5% | 18.5% |
| Mount | 263 | 49 | 488 | 3944 | 1789 | 356 | 21 | 57.1% | 42.9% |
| Shallow Thrust | 378 | 38 | 417 | 1192 | 11208 | 2979 | 16 | 69.1% | 30.9% |
| Deep Thrust | 235 | 3 | 178 | 144 | 1615 | 33384 | 73 | 93.7% | 6.3% |
| Ejaculate | 41 | 15 | 19 | 20 | 37 | 259 | 2092 | 84.3% | 15.7% |

| | Investigate M | Attack | Investigate F | Mount | Shallow Thrust | Deep Thrust | Ejaculate |
|---|---|---|---|---|---|---|---|
| Recall | 85.6% | 89.0% | 85.2% | 63.1% | 70.7% | 87.1% | 91.5% |
| FNR | 14.4% | 11.0% | 14.8% | 36.9% | 29.3% | 12.9% | 8.5% |

**Figure S5. Linear discriminant analysis of recordings associated with specified social behaviors. Related to Figure 5.**

**A.** Distribution of 1st and 2nd LDA components associated with different behaviors from a representative recording session. Each dot represents one data point (frame). The discriminant analysis was performed using only periods containing specified social behaviors.

**B**. Confusion matrix shows the number of frames that are correctly and incorrectly classified for each behavior across all sessions. Left columns show the precision (blue) and false discovery rate (orange). Bottom rows show the recall (blue) and false negative rate (orange).

**C.** F1 scores for various behaviors computed using full models that include data from all recording regions (red) and models built with shuffled data (black). Error bar: mean ± SEM. Inv: investigate; S Thrust: shallow thrust; D Thrust: deep thrust. Paired t-test (if pass Lilliefors normality test) or Wilcoxon signed-rank test (if not pass Lilliefors normality test). All p values are adjusted with Benjamini Hochberg procedure for controlling the false discovery rate. ***$p < 0.001$. $n = 17\text{-}24$ animals. Gray circles represent individual animal results using the full model. See Table S1 for raw data and detailed statistics.

### Figure S6

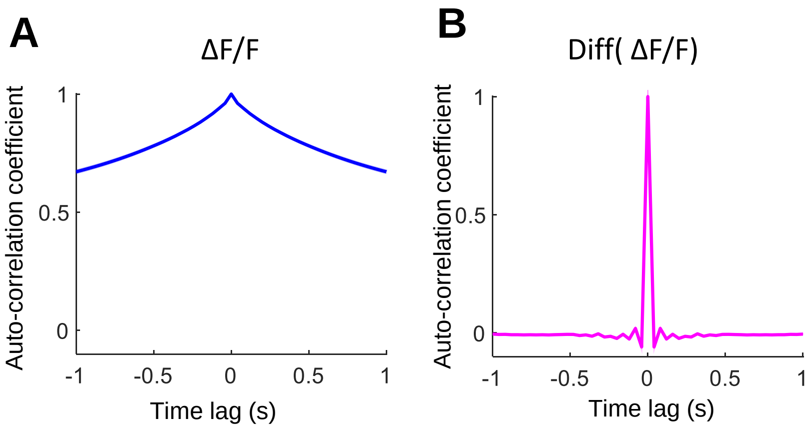


**Figure S6. Auto-correlation plots of Z scored ΔF/F (A) and differential Z scored ΔF/F (B). Related to Figure 6.** The trace is first averaged across all recording regions for one session and then averaged across all sessions. n = 64 sessions. Shades indicate ± SEM.

### Figure S7

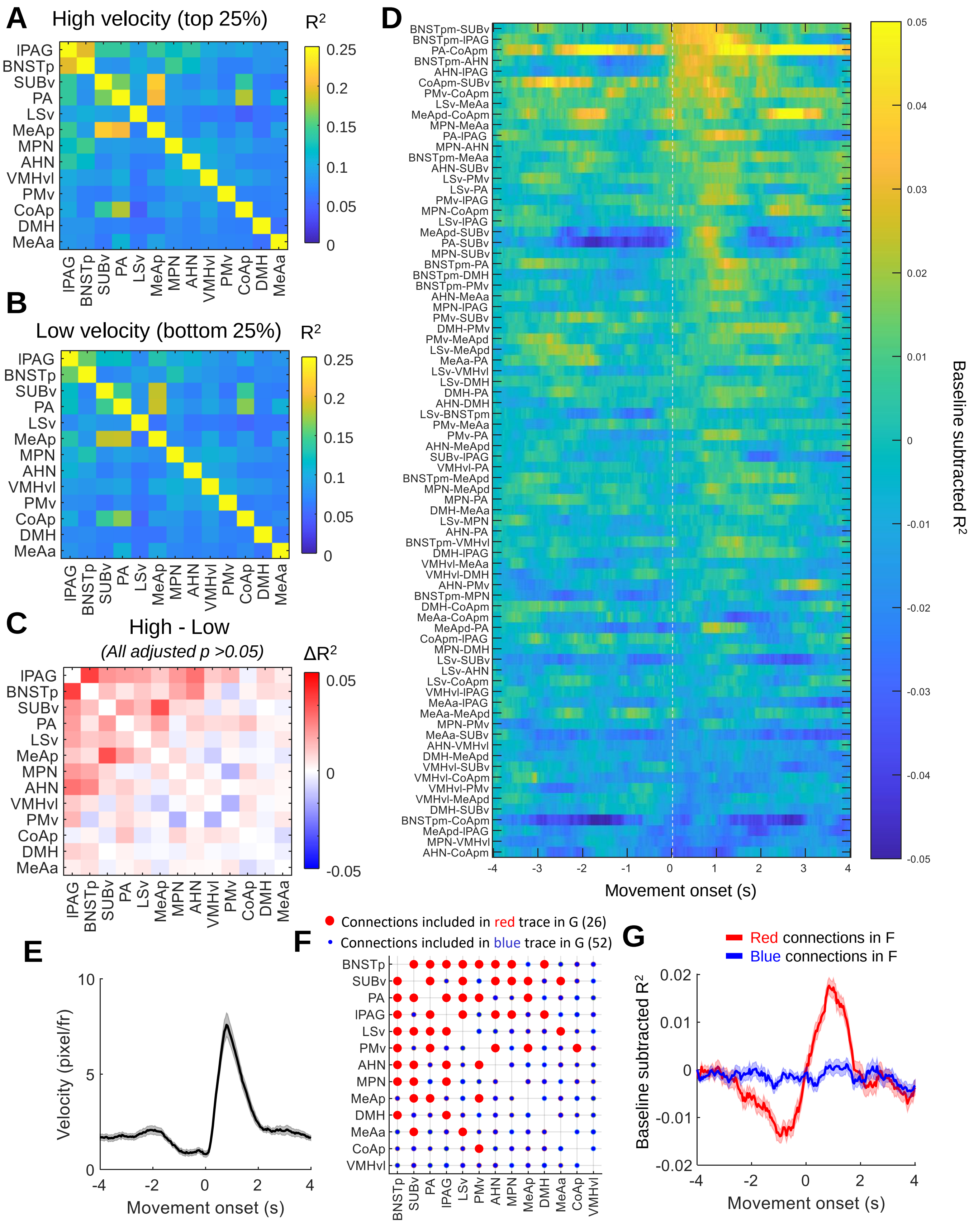
A
High velocity (top 25%)
R²
0.25
0.2
0.15
0.1
0.05
0
IPAG
BNSTp
SUBv
PA
LSv
MeAp
MPN
AHN
VMHvl
PMv
CoAp
DMH
MeAa
B
Low velocity (bottom 25%)
C
High - Low
(All adjusted p >0.05)
ΔR²
0.05
0
-0.05
D
BNSTpm-SUBv
BNSTpm-IPAG
PA-CoApm
BNSTpm-AHN
AHN-IPAG
CoApm-SUBv
PMv-CoApm
LSv-MeAa
MeApd-CoApm
MPN-MeAa
PA-IPAG
MPN-AHN
BNSTpm-MeAa
AHN-SUBv
LSv-PMv
LSv-PA
PMv-IPAG
MPN-CoApm
LSv-IPAG
MeApd-SUBv
PA-SUBv
MPN-SUBv
BNSTpm-PA
BNSTpm-DMH
BNSTpm-PMv
AHN-MeAa
MPN-IPAG
PMv-SUBv
DMH-PMv
PMv-MeApd
LSv-MeApd
MeAa-PA
LSv-VMHvl
LSv-DMH
DMH-PA
AHN-DMH
LSv-BNSTpm
PMv-MeAa
PMv-PA
AHN-MeApd
SUBv-IPAG
VMHvl-PA
BNSTpm-MeApd
MPN-MeApd
MPN-PA
DMH-MeAa
LSv-MPN
AHN-PA
BNSTpm-VMHvl
DMH-IPAG
VMHvl-MeAa
VMHvl-DMH
AHN-PMv
BNSTpm-MPN
DMH-CoApm
MeAa-CoApm
MeApd-PA
CoApm-IPAG
MPN-DMH
LSv-SUBv
LSv-AHN
LSv-CoApm
VMHvl-IPAG
MeAa-IPAG
MeAa-MeApd
MPN-PMv
MeAa-SUBv
AHN-VMHvl
DMH-MeApd
VMHvl-SUBv
VMHvl-CoApm
VMHvl-PMv
VMHvl-MeApd
DMH-SUBv
BNSTpm-CoApm
MeApd-IPAG
MPN-VMHvl
AHN-CoApm
-4
-3
-2
-1
0
1
2
3
4
Movement onset (s)
0.05
0.04
0.03
0.02
0.01
0
-0.01
-0.02
-0.03
-0.04
-0.05
Baseline subtracted R²
E
Velocity (pixel/fr)
10
5
0
-4
-2
0
2
4
Movement onset (s)
F
Connections included in red trace in G (26)
Connections included in blue trace in G (52)
BNSTp
SUBv
PA
IPAG
LSv
PMv
AHN
MPN
MeAp
DMH
MeAa
CoAp
VMHvl
G
Red connections in F
Blue connections in F
0.02
0.01
0
-0.01
-0.02
Baseline subtracted R²
Movement onset (s)

**Figure S7. Change in functional connectivity in the expanded SBN with movement. Related to Figure 7.**

**A and B.** Average $R^2$ of each connection (a pair of regions) during the high-velocity period (top 25%) **(A),** and low-velocity period (bottom 25%) **(B),** when the recording animal was alone in its home cage.

**C.** The difference in R2 between the high-velocity period and low-velocity period. Paired t-test for each connection. All p values are adjusted with Benjamini Hochberg procedure for controlling the false discovery rate.

**D.** Heat maps showing the change in $R^2$ aligned to the movement onset. The R2 values are corrected by subtracting the value at -4s for each connection. The plots are sorted based on the mean baseline-corrected R2 value between 0-1s after the movement onset.

**E.** Animal's velocity aligned to the onset of the movement. n = 25 animals. Shades: ± SEM.

**F.** Movement-sensitive connections (red circles, $\Delta R^2>0.01$) and -insensitive connections (blue dots, $\Delta R^2>0.01$). $\Delta R^2$ is calculated as the difference in $R^2$ between 0-1s after movement onset and -1-0s before movement onset.

**G.** The average baseline subtracted R2 of movement-sensitive (red, n = 26) and insensitive (blue, n =52) connections. Shades: ± SEM.

See Table S1 for raw data and detailed statistics.

### Figure S8

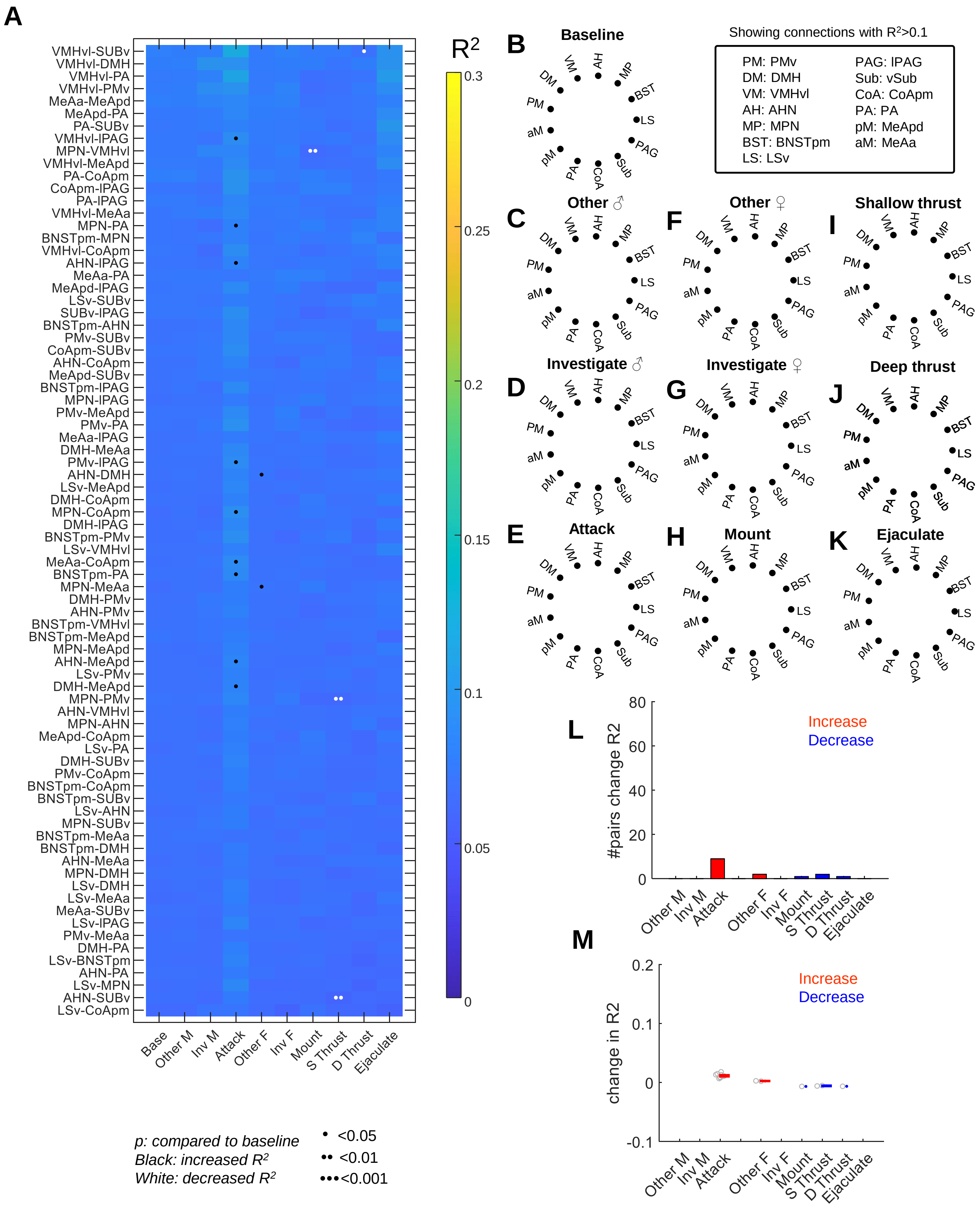
A
VMHvl-SUBv
VMHvl-DMH
VMHvl-PA
VMHvl-PMv
MeAa-MeApd
MeApd-PA
PA-SUBv
VMHvl-lPAG
MPN-VMHvl
VMHvl-MeApd
PA-CoApm
CoApm-lPAG
PA-lPAG
VMHvl-MeAa
MPN-PA
BNSTpm-MPN
VMHvl-CoApm
AHN-lPAG
MeAa-PA
MeApd-lPAG
LSv-SUBv
SUBv-lPAG
BNSTpm-AHN
PMv-SUBv
CoApm-SUBv
AHN-CoApm
MeApd-SUBv
BNSTpm-lPAG
MPN-lPAG
PMv-MeApd
PMv-PA
MeAa-lPAG
DMH-MeAa
PMv-lPAG
AHN-DMH
LSv-MeApd
DMH-CoApm
MPN-CoApm
DMH-lPAG
BNSTpm-PMv
LSv-VMHvl
MeAa-CoApm
BNSTpm-PA
MPN-MeAa
DMH-PMv
AHN-PMv
BNSTpm-VMHvl
BNSTpm-MeApd
MPN-MeApd
AHN-MeApd
LSv-PMv
DMH-MeApd
MPN-PMv
AHN-VMHvl
MPN-AHN
MeApd-CoApm
LSv-PA
DMH-SUBv
PMv-CoApm
BNSTpm-CoApm
BNSTpm-SUBv
LSv-AHN
MPN-SUBv
BNSTpm-MeAa
BNSTpm-DMH
AHN-MeAa
MPN-DMH
LSv-DMH
LSv-MeAa
MeAa-SUBv
LSv-lPAG
PMv-MeAa
DMH-PA
LSv-BNSTpm
AHN-PA
LSv-MPN
AHN-SUBv
LSv-CoApm
Base
Other M
Inv M
Attack
Other F
Inv F
Mount
S Thrust
D Thrust
Ejaculate
R2
0.3
0.25
0.2
0.15
0.1
0.05
0
p: compared to baseline
Black: increased R2
White: decreased R2
<0.05
<0.01
<0.001
B Baseline
Showing connections with R2>0.1
PM: PMv
DM: DMH
VM: VMHvl
AH: AHN
MP: MPN
BST: BNSTpm
LS: LSv
PAG: lPAG
Sub: vSub
CoA: CoApm
PA: PA
pM: MeApd
aM: MeAa
C Other ♂
D Investigate ♂
E Attack
F Other ♀
G Investigate ♀
H Mount
I Shallow thrust
J Deep thrust
K Ejaculate
DM VM AH MP BST LS PAG Sub CoA PA pM aM PM
L
#pairs change R2
80
60
40
20
0
Increase
Decrease
Other M
Inv M
Attack
Other F
Inv F
Mount
S Thrust
D Thrust
Ejaculate
M
change in R2
0.2
0.1
0
-0.1
Increase
Decrease
Other M
Inv M
Attack
Other F
Inv F
Mount
S Thrust
D Thrust
Ejaculate

**Figure S8. Correlation across regions is abolished after slightly jittering the simultaneously recorded traces. Related to Figure 7.**

**A.** After adding a small random jitter (40-200ms) to the simultaneously recorded traces, the average $R^2$ values of all pairs of regions during all behavior epochs are close to 0 and largely not different from that during the pre-intruder baseline period. “Base” refers to pre-intruder period. “Other M” and “Other F” refer to periods when the male or female intruder is present but no specific social behavior is annotated. Inv: investigate; S Thrust: shallow thrust; D Thrust: deep thrust. Paired t-test (if pass Lilliefors normality test) or Wilcoxon signed-rank test (if not pass Lilliefors normality test). P values are adjusted using with Benjamini Hochberg procedure for controlling the false discovery rate. *$p<0.05$; **$p<0.01$; ***$p<0.001$. Black and white indicate a significant increase or decrease from the baseline, respectively.

**B-K.** Graph plots showing the strength of functional connectivity ($R^2$) among different regions during various social behavior epochs using jittered recording data. No connection showed $R^2 > 0.1$ during any behavior.

**L.** the number of pairs of regions that show significantly increased $R^2$ (red) or decreased $R^2$ (blue) from the pre-intruder baseline using jittered traces.

**N.** Change in R2 values from the pre-intruder baseline for significantly changed connections using jittered traces. Red and blue show the mean ± SEM of significantly increased and decreased connections during each behavior. n = 0-9 pairs of regions.

Note that as the jitter is random, the results vary slightly for each run. See Table S1 for raw data and detailed statistics.

### Figure S9

| ♂ intruder | | MPN | AHN | VMHvl | DMH | PMv | BNSTp | MeAa | MeAp | PA | CoAp | SUBv | lPAG | LSv | FC |
|---|---|---|---|---|---|---|---|---|---|---|---|---|---|---|---|
| Initial encounter | Largest | | | ● | | | | | | | | | | | |
| | Fastest | | | | ● | | | | | | | | | | |
| Investigate | Largest | | | ● | | | | | | | | | | | ↑ |
| | Fastest | | | | | | | ● | | | | | | | |
| Attack | Largest | | | ● | ● | | | | | | | | ● | | ↑↑↑ |
| | Fastest | | | ● | | | | | | | | | | | |

| ♀ intruder | | MPN | AHN | VMHvl | DMH | PMv | BNSTp | MeAa | MeAp | PA | CoAp | SUBv | lPAG | LSv | FC |
|---|---|---|---|---|---|---|---|---|---|---|---|---|---|---|---|
| Initial encounter | Largest | ● | | | | | | | | | | | | | |
| | Fastest | | | | ● | | | | | | | | | | |
| Investigate | Largest | ● | | | | | | ● | | ● | | | | | ↑ |
| | Fastest | | | | | | | ● | | | | | | | |
| Mount | Largest | ● | | | | | | ● | | | | | | | ↑↑ |
| | Fastest | | | | | | | ● | | | | | | | |
| S Thrust | Largest | ● | | | | | | | | | | | | | ↔ |
| D Thrust | Largest | ● | | | | | | | | | | | | | ↓ |
| Ejaculate | Largest | | | | | | ● | | | | | | | | ↓↓ |

**Figure S9.** Summary of the response patterns in the expanded SBN during social behaviors in male mice. FC: functional connectivity.
